# Cell density-dependent death triggered by viral palindromic DNA sequences

**DOI:** 10.1101/2022.11.18.517080

**Authors:** William P Robins, Bradley T Meader, Jonida Toska, John J Mekalanos

## Abstract

Defense systems that recognize viruses have provided not only the tools of molecular biology but also important insights into other mechanisms that can induce immunity to these or other infectious agents including transmissible plasmids and chromosomal genetic elements. Several systems that trigger cell death upon viral infection have recently been recognized but the signals that activate these abortive infection systems remain largely unknown. Here we characterize one such system in *Vibrio cholerae* that we found was responsible for triggering <u>c</u>ell-density <u>d</u>ependent <u>d</u>eath (CDD) of bacterial cells in response to the presence of certain genetic elements. The key components of the CDD system include quorum-regulated components DdmABC and the host factor PriA. Our analysis indicates that the plasmid and phage signals that trigger CDD were palindromic DNA sequences that are predicted to form stem-loop hairpin structures from single-stranded DNA during stalled replication. Our results further suggest that other agents that generate damaged DNA can also trigger DdmABC/PriA activation and cell death probably through activation of a nuclease domain present in the DdmA protein. Thus, any infectious process that results in damaged DNA, particularly during DNA replication, can in theory trigger cell death through the DdmABC/PriA abortive infection system.

## Introduction

Anti-viral defense systems have a long history of identification in procaryotic systems through both bacteriophage and bacterial genetics ^1^. Many of these antiphage systems have been recently recognized to have eukaryotic counterparts that function in viral defense ^2,3^. Recent advances in genomics and bioinformatics have facilitated the prediction of numerous putative antiphage systems that have uncharacterized mechanisms of action ^4^. The genes for these antiphage systems are often found positioned together on mobile genetic elements that were likely horizontally transferred between bacterial strains ^5^. Frequently, these anti-phage systems have been experimentally shown to function in *Escherichia coli* ^6–8^, however, the signals that allow such systems to recognize phage-infected bacterial cells remain ill-defined.

Phages are the most abundant biological entities on planet Earth and are predicted to outnumber bacteria by a factor of 10-fold ^9^. Given this continuous predation, bacteria have evolved antiphage systems and, in turn, phages have evolved escape strategies for these formidable defenses. Together these define an enormous ‘arms race’ that shapes microbial ecology in all inhabited environments ^10–12^. It is important to note that understanding how anti-phage defense systems work at a mechanistic level has provided numerous technological tools that now shape and drive biomedical and bioscience innovations ^13,14^

Phage and mobile elements in bacterial pathogens are also highly relevant to diseases that include the lethal diarrheal syndrome called cholera ^15,16^. For pathogenic *Vibrio cholerae*, the acquisition and chromosomal integration of the genome of the filamentous phage (CTX-ϕ encoding cholera toxin was clearly a prerequisite for the emergence of strains responsible for all seven recognized cholera pandemics ^17^. However, the process that leads to the acquisition of CTX-ϕ involves multiple phage-like elements and a chromosomal island that encodes the Toxin-Co-regulated Pilus (TCP) which serves as both the receptor for the phage as well as a critical human intestinal colonization factor ^17–20^.

All pandemic strains of *V. cholerae* are genetically similar and classified as either Classical or El Tor biotypes ^21^. Classical strains are believed to be responsible for the first six pandemics and separate waves of El Tors variants account for the current ongoing 7th pandemic ^15,16,22–24^. Other genetically distinct strains have acquired both TCP and CTX-ϕ but these strains are not responsible for widespread epidemics ^25–27^. Thus, the unique repertoire of genetic elements in the 7th pandemic El Tor strains is believed to contribute to the fitness of this dominant pandemic clade of *V. cholerae* that emerged with the subsequent elimination of Classical strains as the cause of cholera worldwide ^16,28^.

*V. cholerae* has been documented to be subject to phage predation that correlates with the collapse of cholera epidemics and this observation is postulated to be a factor in the emergence of the 7th pandemic clade ^29–32^. These observations have prompted many studies on phage resistance mechanisms in *V. cholerae*, as well as phage-encoded countermeasures that block some anti-phage systems ^33,34^. For example, quorum sensing regulation mediated by autoinducers, molecules that accumulate typically under high-density growth conditions, was reported to promote phage resistance in 7th pandemic *V. cholerae* strains by an unclear mechanism ^35^.

All contemporary pathogenic *V. cholerae* strains have acquired two chromosomal islands termed the *Vibrio* 7th Pandemic Islands I (VSP-1) and II (VSP-2) ^28^. Gene products encoded by both islands have recently been shown to be capable of restricting phage growth and plasmid persistence by mechanisms including programmed cell death of infected cells ^7,6,8,36^. These conclusions have been largely reached based on the over-expression of VSP-1 or VSP-2 genes in the surrogate bacterial species *E*. coli. More recently, quorum-regulated VSP-2 genes in the *DdmABC* operon have been demonstrated to inhibit the growth of some vibriophages by 2-3 logs but only when co-absent with specific VSP-1 putative antiphage genes ^36^. Importantly, the signals that activate any anti-phage and anti-plasmid defense system in *V. cholerae* remain ill-defined.

To discover the signals that activate these defense systems in *V. cholerae*, we analyzed a laboratory plasmid and two phages we identified that were specifically restricted in growth by VSP-2 genes *DdmABC* that belong to the Lamassu-like antiphage system ^4^. When maintained by drug selection, the transformed plasmid remarkably caused bacterial death only at high cell density and in a VSP-2-dependent manner. Using genetic approaches, we identified the cell-death-inducing signal as a sequence in the plasmid that is predicted to form a single-stranded DNA stem loop (or hairpin). We cloned sequences from vibriophages that were similarly toxic and dependent on VSP-2 and found that all were palindromic and predicted to form stem-loop hairpin structures. Additional VSP-2 mutations that could suppress the high-density dependent, hairpin-triggered cell death response mapped to the quorum-regulated *DdmABC* operon as well as a gene product (PriA) that recognizes stalled DNA replication forks. DNA-damaging agents also triggered higher levels of cell death dependent on VSP-2, DdmABC, and PriA function. We present a model for how phage infection is detected by the DdmABC/PriA system and how that recognition likely triggers cell death through the activation of a nuclease domain encoded by the DdmA protein. If correct, this model predicts that the DdmABC/PriA system can detect not only phages and plasmids that encode palindromes which likely damage DNA during replication but also viral and host processes that cause chromosomal or viral DNA degradation.

## Results

### Identification of a sequence in a plasmid that triggers *V. cholerae* cell death

El Tor 7th pandemic strains of *V. cholerae* have been reported to inhibit the maintenance of certain plasmids ^8^. To independently confirm this observation, we attempted to transform a plasmid pWR1566 that encodes LacZ and ampicillin (Amp) resistance into a *lacZ* mutant of a 7th pandemic El Tor *V. cholerae* strain (WPR2700). LacZ expression was monitored with the colorimetric substrate Bromo-4-Chloro-3-Indolyl-β-d-Galactopyranoside (XGAL) which forms a dark blue product when hydrolyzed by LacZ. Although LacZ expression was observed in Amp-selected transformants, colonies grew poorly and released copious amounts of LacZ as indicated by the appearance of the dark blue halos around colonies. These results suggested that cell death and lysis were occurring; a phenomenon previously described as “blue ghosts” in *E. coli* ^37^ (**Figure 1a)**. Surprisingly, WPR2700 cells carrying the plasmid grew well in liquid media containing Amp, but when those high-density cultures were plated on agar containing Amp, we observed cell death that was most apparent where cells were present at high density; accordingly, we called this phenomenon, <u>c</u>ell density-<u>d</u>ependent <u>d</u>eath (CDD) (**Figure 1b**).

**Figure 1.**
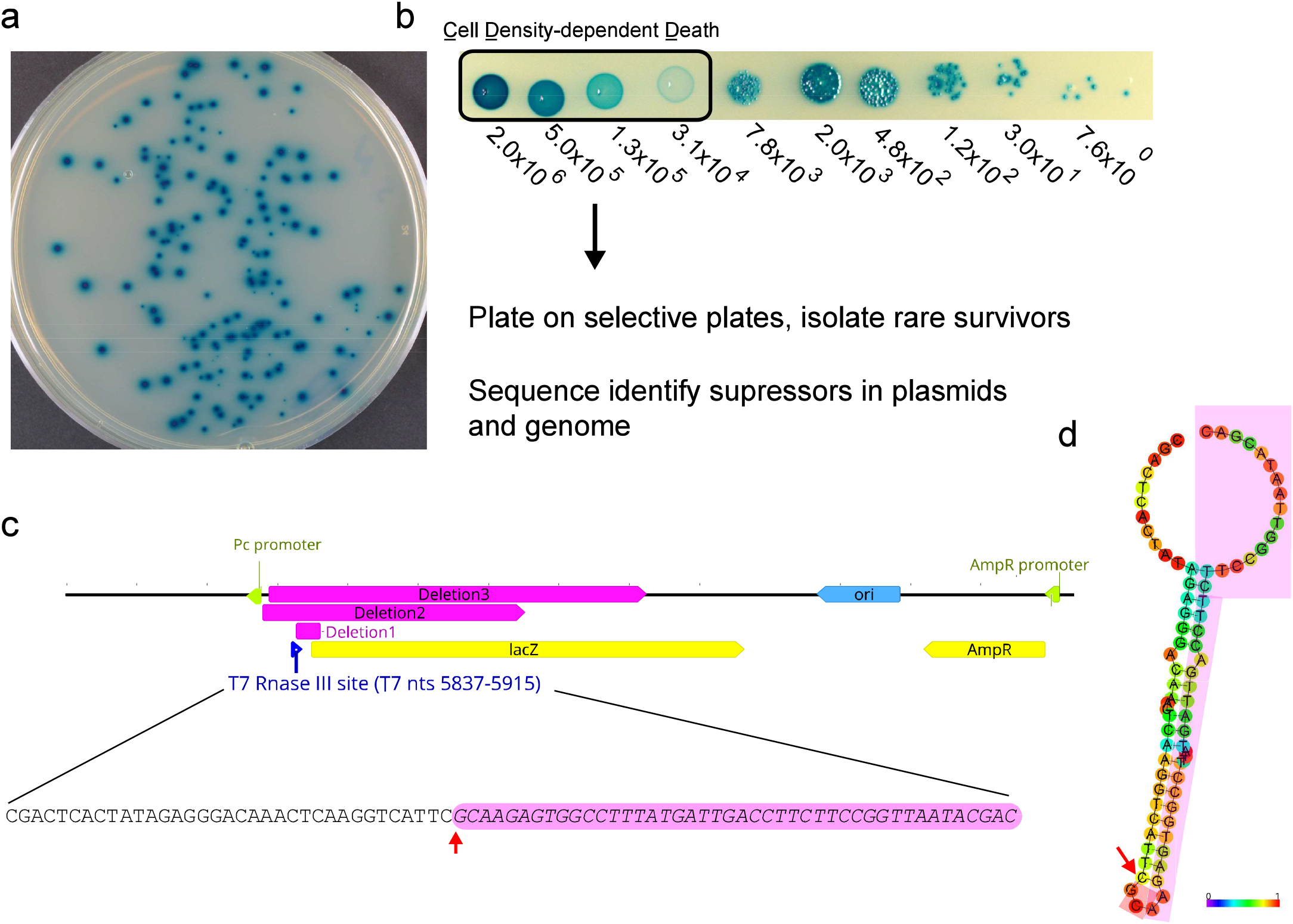
CCD phenotype and selection for suppressors reveal a *cis-*acting sequence in a plasmid. a| Blue ghost phenotype when transformed cells grow as colonizes and release b-galactosidase. b| A cell-dependent death (CDD) phenotype is observed when a LacZ-containing plasmid is transformed into *V*.*cholerae* and the transformants are plated at high density under plasmid-selecting conditions. c| Map of the plasmid showing deletions (in pink) identified in recovered plasmid mutants that enable survival. d| Sequence of T7 RNAse-III site common within all deletions and predicted secondary structure. A colorimetric confidence scale for base pairing prediction is shown.

Because CDD occurred only at high cell density on a solid surface, we reasoned this may be due to quorum sensing as well as recognition of a signal present on the plasmid that triggered cell death. To isolate mutants resistant to the CDD signal, plasmid-transformed WPR2700 cells were spotted at high density on agar containing Amp and XGAL. While most cells died, we did observe exceedingly rare white colonies that arose at a frequency of approximately 10^−7^. Because the majority of these Amp^r^ clones no longer produced LacZ, we predicted that plasmid sequence deletions had occurred and indeed sequencing of the purified mutant plasmids revealed deletions that removed a common 5’ region upstream of the *lacZ* gene (**Figure 1c**). Curiously, the formation of these deletions had not been generated between short homologous sequences as previously observed as the most common mechanism for deletion formation ^38^ (**Supplemental Figure 1**). We reasoned that this deleted region must encode a *cis*-acting sequence that induces CDD. Examination of deleted sequences revealed a 79 nucleotide plasmid sequence originally derived from the primary replication origin of *E. coli* bacteriophage T7 including an intact annotated RNAse-III site (**Figure 1cd)**. These sequences form hairpins as dsRNA and then are processed by RNases to regulate transcript stability and gene expression ^39^. There is no known role for these secondary structures in DNA, though they are energetically favored to form in single-stranded nucleic acid sequences including DNA because of their palindromic nature. Plasmids lacking this sequence due to a deletion or related plasmids that lacked this same T7-derived sequence but carried LacZ did not induce CDD.

### Identification of *V. cholerae* suppressors of CDD

Using the same selective approach, we identified *V. cholerae* chromosomal mutants that could maintain and replicate pWPR1566 and grow as healthy blue colonies on agar containing Amp and XGAL. Next-generation sequencing of each of these mutants was used to identify CDD suppressor alleles in the *V. cholerae* chromosomal genome and also estimate the copy number of pWR1566 through the level of reads that mapped to the plasmid. Of 24 mapped suppressor mutations in blue colonies that were chromosomal, five were found to affect plasmid copy number based on mapping depth (**Table 1**). All five mutations decreased plasmid copy 8-20 fold and four mutations were located in the gene encoding PcnB (PAP-I), a protein known to decrease the copy number of plasmids with a pCol-E1 type replicon, including our vector (Lopilato et al., 1986; He et al., 1993); the fifth mutation was in *priA*, a gene encoding the PriA primosome and replication restart protein also essential for replication of pCol-E1 plasmids ^40^. This result suggested the reduction in plasmid-encoded signal permitted survival. A majority (18 of 24) of the suppressor mutations mapped to three adjacent genes in VSP-2 island that encode the *DdmABC* operon ^8^. These include missense, nonsense, frameshifts, deletions, and insertions. Thus, we speculated that the DdmC proteins were part of the DDC system responsible for the plasmid-induced cell death that was triggered by growth at high cell density. Other suppressor alleles independent of *DdmABC* and *PcnB* included a mutation in *luxO* (**Table 1**); the *luxO* mutation is consistent with the observation that the DdMABC operon is quorum-regulated ^36^.

**Table 1.**
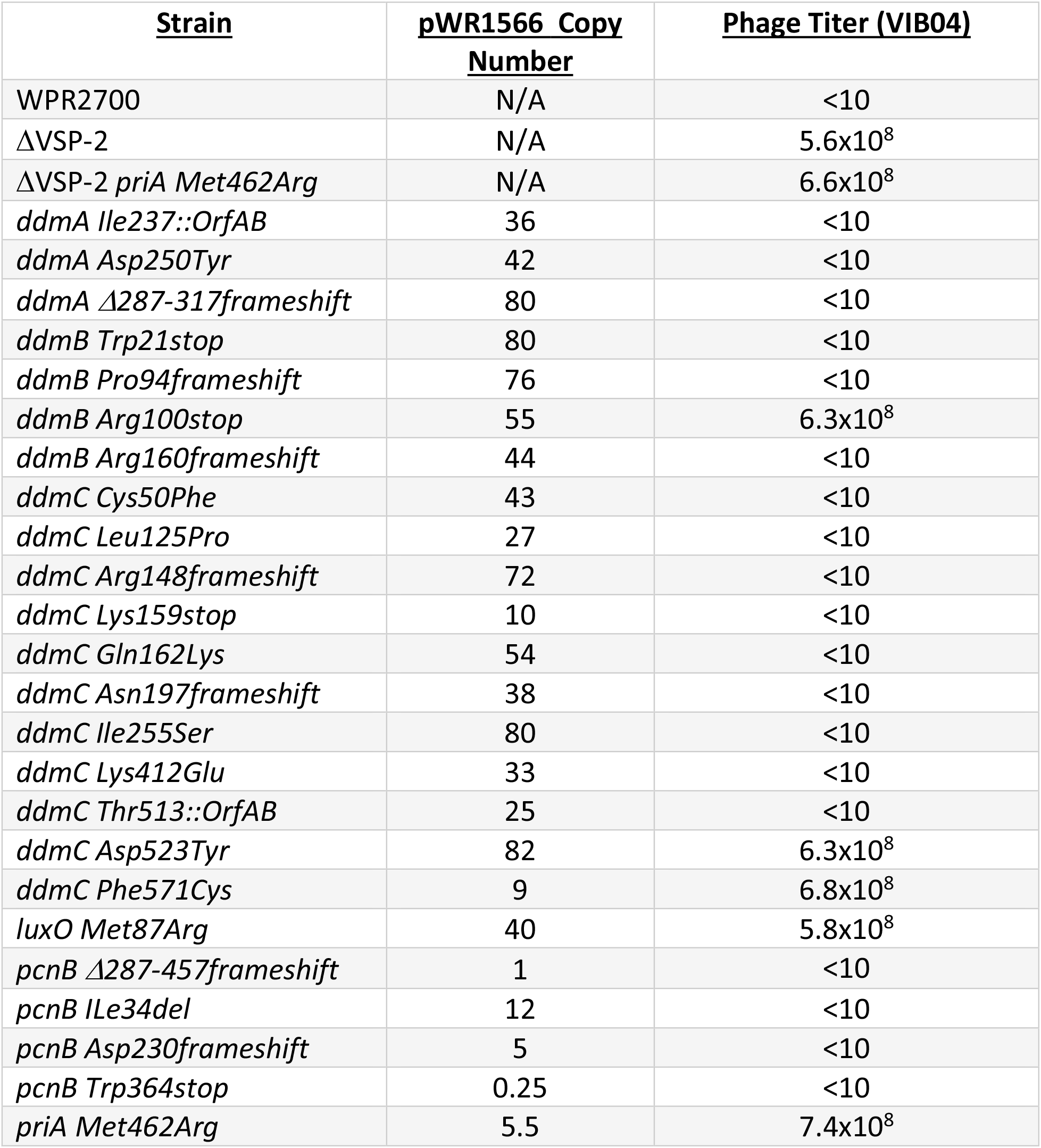
Identified genomic suppressors and their effects on plasmid copy number and VIB04 phage titers. In addition to VSP-2 and non-VSP-2 mutants, WR2700 and WR2700 *ΔVSP-2* are included as controls permissive and restrictive for phage growth.

| **<u>Strain</u>** | **<u>pWR1566 Copy Number</u>** | **<u>Phage Titer (VIB04)</u>** |
| --- | --- | --- |
| WPR2700 | N/A | <10 |
| ΔVSP-2 | N/A | $5.6 \times 10^8$ |
| ΔVSP-2 *priA* Met462Arg | N/A | $6.6 \times 10^8$ |
| *ddmA* Ile237::*OrfAB* | 36 | <10 |
| *ddmA* Asp250Tyr | 42 | <10 |
| *ddmA* Δ287-317frameshift | 80 | <10 |
| *ddmB* Trp21stop | 80 | <10 |
| *ddmB* Pro94frameshift | 76 | <10 |
| *ddmB* Arg100stop | 55 | $6.3 \times 10^8$ |
| *ddmB* Arg160frameshift | 44 | <10 |
| *ddmC* Cys50Phe | 43 | <10 |
| *ddmC* Leu125Pro | 27 | <10 |
| *ddmC* Arg148frameshift | 72 | <10 |
| *ddmC* Lys159stop | 10 | <10 |
| *ddmC* Gln162Lys | 54 | <10 |
| *ddmC* Asn197frameshift | 38 | <10 |
| *ddmC* Ile255Ser | 80 | <10 |
| *ddmC* Lys412Glu | 33 | <10 |
| *ddmC* Thr513::*OrfAB* | 25 | <10 |
| *ddmC* Asp523Tyr | 82 | $6.3 \times 10^8$ |
| *ddmC* Phe571Cys | 9 | $6.8 \times 10^8$ |
| *luxO* Met87Arg | 40 | $5.8 \times 10^8$ |
| *pcnB* Δ287-457frameshift | 1 | <10 |
| *pcnB* Ile34del | 12 | <10 |
| *pcnB* Asp230frameshift | 5 | <10 |
| *pcnB* Trp364stop | 0.25 | <10 |
| *priA* Met462Arg | 5.5 | $7.4 \times 10^8$ |

### Predicted structures and functions of VSP-II proteins

By homology alone, DdmABC belongs to the Lamassu antiphage system (**Figure 2cd**) ^4^. Hidden Markov Model and both our and previous *in silico* structural analyses identified a conserved CAP4 nuclease domain in DdmA that is also present in a CBASS antiphage nuclease effector protein that requires a cyclic tri oligonucleotide for its activation by binding a domain called SAVED ^8,36,41^; however, DdmA does not have a SAVED domain but instead a C-terminal Domain 7 motif (Pfam CDT7) belonging to ABC-three component (ABC-3C) systems. DdmB is predicted to be a small globular protein with no predicted homologs with known functions but does possess an ABC-three component system Middle Component motif (Pfam MC3). We detected a strong predicted structural similarity between DdmC and the eukaryotic Rad50 and bacterial SbcC-like proteins (**Supplemental Figure 2**). Features found in other Rad50/SbcC-like proteins (a conserved Walker A and an atypical Walker B NTP-binding motif) were independently verified as in previous work ^8^, and identified in each of the head domains at the N- and C-termini of DdmC, respectively. The three proteins encoded by the *DdmABC* operon closely resemble other members of the recently identified ABC-3C (ABC-three component) complex (**Figure 2e**). These are defined by a toxic effector enzyme that attacks an essential cellular process when partner proteins in the complex recognize signals such as those presented by an invading DNA virus ^42^.

**Figure 2.**
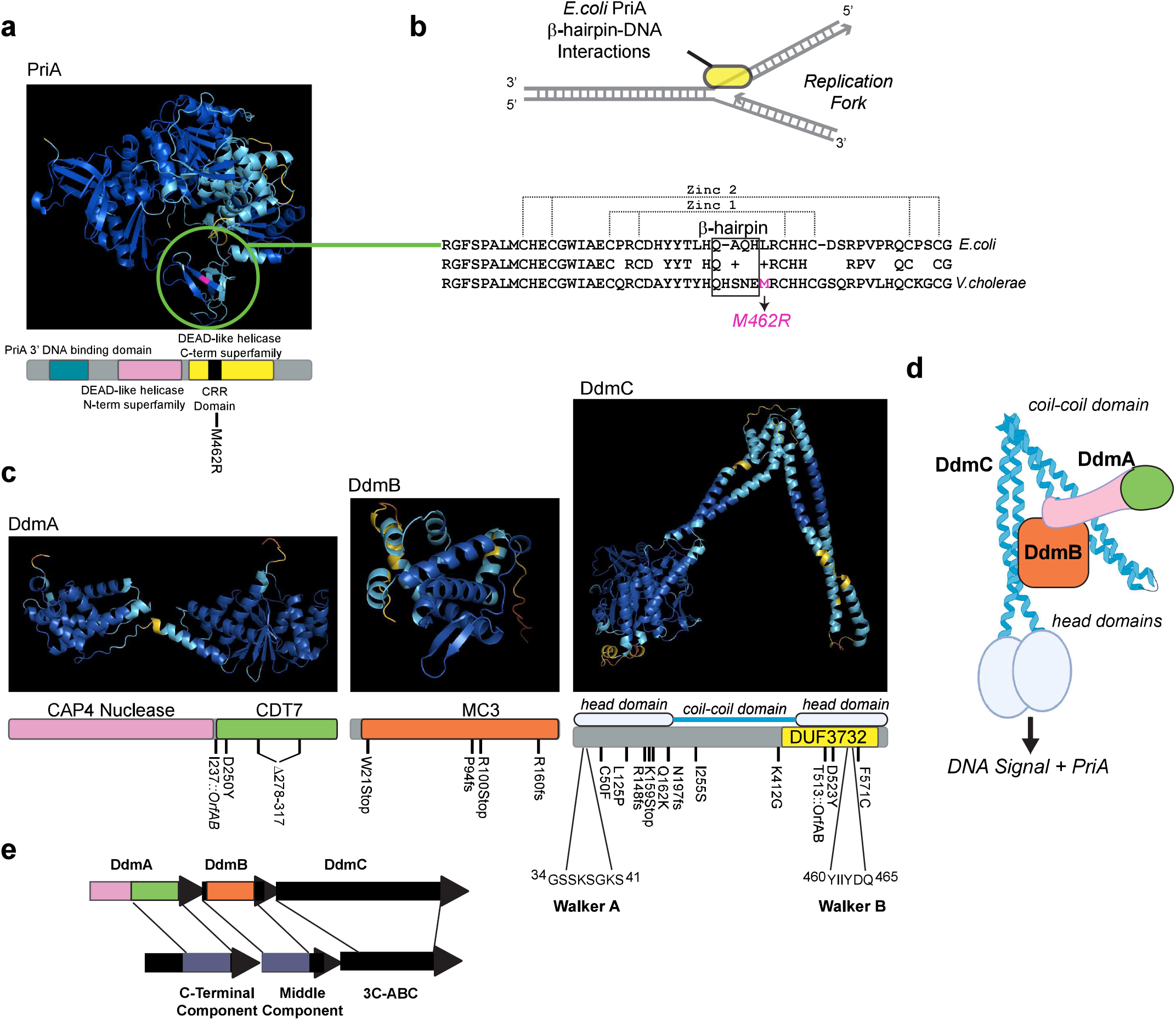
Colabfold predictions of *V*.*cholerae* PriA and the three proteins encoded in the DdmABC operon. a|Predicted structure of *V*.*cholerae* PriA and the represented domains conserved in other bacterial PriA proteins. The CRR domain possesses the M462R mutation identified in this work. The CRR/b-hairpin domain is circled. b| Alignment of *V*.*cholerae* and *E*.*coli* b-hairpin domain residues. The CRR domain is boxed and the position of the M462R is indicated. Nucleotides known to interact with the *E*.*coli* PriA β-hairpin in a stalled replication fork are shown. c|Predicted PDB structures of DdmA, DdmB, and DdmC and predicted domains and motifs. Mutations identified in this work are indicated. d| Cartoon model showing a general DdmABC complex and what this may identify during restriction of phage or plasmids. e|Homology between DdmABC and 3C-ABC complexes based on conserved domains present in proteins.

We mapped the CDD suppressor mutations in the *DdmABC* operon and compared their positions to the predicted structures using ColabFold ^43^. All mutations in *ddmA* were either insertion, missense, deletions, frameshift, or premature stop codons indicating that DdmA loss of function was the most likely cause of CDD suppression. All *ddmB* mutations were either frameshift or stop codons also suggesting loss of function. For *ddmC*, most mutations were clustered in the two predicted head domains and included premature stop codons, frameshift, and missense mutations. The DdmC F571C mutation was notably proximal to the predicted Walker B motif at AA 560-565. Because of the limited scope of this genetic selection, we did not expect to identify precise mutations in all domains essential for DdmABC activity such as key residues within the DdmA nuclease domain.

### Identification of VSP-2 restricted phages and phage-associated signals that trigger CDD

We further reasoned that CDD may represent a response usually associated with aborted phage infection triggered by DNA sequences such as the T7 phage-derived hairpin/T7 Rnase-III site we identified in pWR1566 and that these sites might be recognized by DdmABC and perhaps PriA if such sequences also stall DNA replication processes. To test this hypothesis, we screened 48 vibriophage samples in our collection using both WT and a constructed VSP-2 deletion mutant (ΔVSP-2) to identify those that were restricted by VSP-2. Two vibriophages (VIB04 and VIB05) were identified as being restricted and both plated as clear large plaques at least 10 million fold better on a confluent lawn of the WR2700Δ*VSP-2* strain (**Table 1**). The genome sequence of the two vibriophages revealed that VIB04 is highly similar to coliphage T7/T3-like viruses and VIB05 to the Tawavirus genera. Both are related and taxonomically assigned within the Autographiviridae family with other similar phages that encode a self-transcribing RNA polymerase ^44^. We were unable to isolate spontaneous mutants in either phage that overcame apparent VSP-2 restriction using high titer cesium-purified stocks with concentrations exceeding ∼10^9^ PFU/ml.

We reasoned one or more DNA signals such as palindromic sequences must exist in both these phages that are restricted by VSP-2 and that these may be essential for phage replication. To screen for such sequences in phage, we chose to focus on phage VIB04 and sheared purified DNA to 400-600 nucleotides and constructed random shotgun libraries cloned into our LacZ^+^ screening vector (pWR1566) while removing the T7-based palindromic sequence which we showed to trigger CDD in a VSP-2-dependent fashion. Liquid cultures of *V. cholerae* carrying these libraries were plated and colonies that displayed the ‘blue ghost’ CDD death phenotype were identified in a small fraction of each shotgun library; plasmids recovered from these colonies were then sequenced. Cloned sequences from VIB04 that induced the blue ghost death phenotype were interspersed across the entire genomes. Many possess high similarities to one another and all encode palindromic sequences that would form hairpins in single strands of DNA and some of these were highly similar to the Rnase-III site in T7 originally identified in plasmid pWR1566 **(Figure 3, Supplemental Figure S3**). Furthermore, these phage-encoded palindromic sequences displayed no toxicity when introduced into *ddmABC* or *priA* mutant strains that suppress CDD.

**Figure 3.**
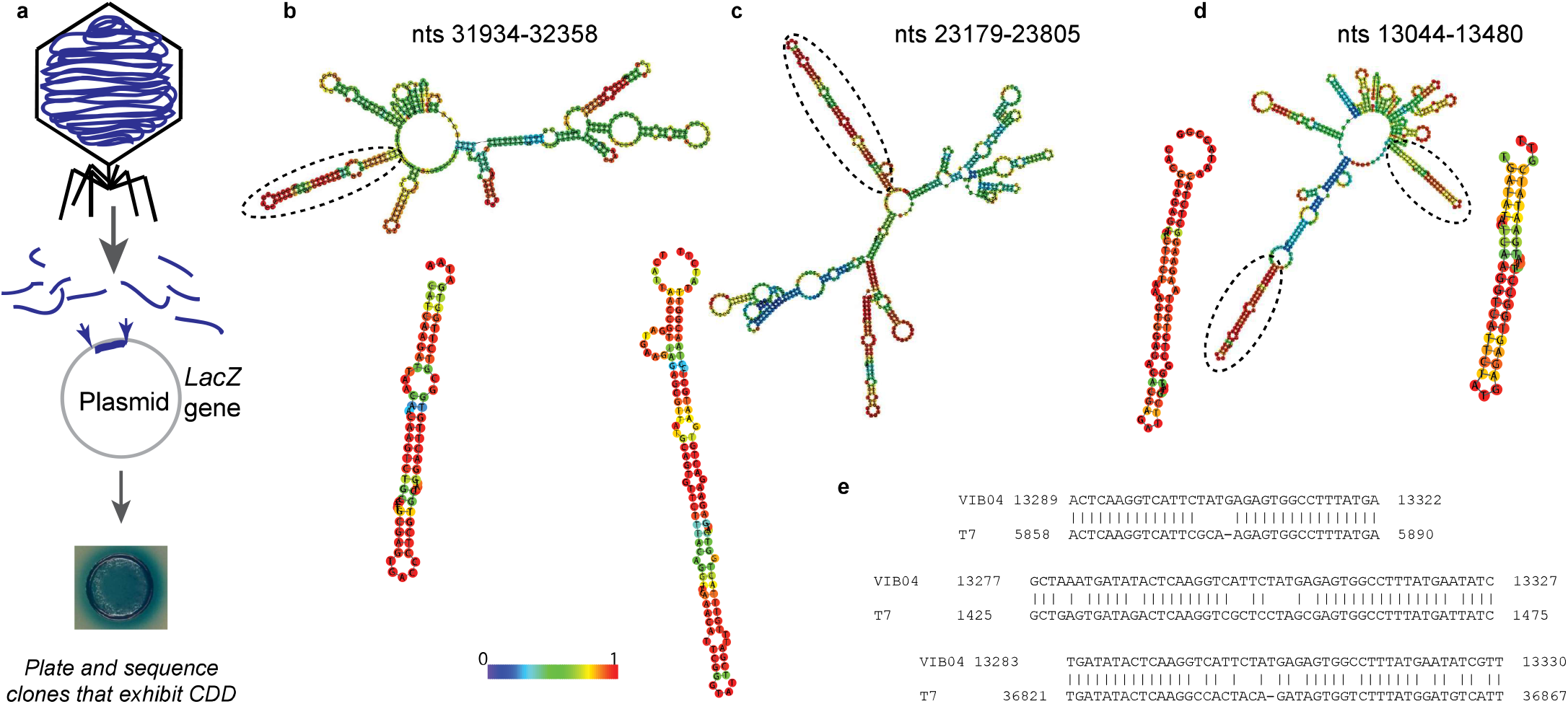
Sequences from restricted phages cloned in place of the T7 Rnase III sequence cause CDD. a | Diagram showing the cloning of random phage DNA fragments into a plasmid and screening for CDD. b-d | Sequences identified in phage VIBO4 by position and predicted secondary structure. A colorimetric confidence scale for base pairing prediction is shown. e | One VIB04 sequence is homologous to the RNaseIII sites in related bacteriophage T7.

### Role of VSP-2 independent suppressors including PriA

We plated both VSP-2 restricted phage VIB04 on the entire set of selected and sequenced *V. cholerae* chromosomal suppressors of CDD to determine if the suppressors obtained by our pWR1566 plasmid selection also permitted phage growth. Some VSP-2 and non-VSP-2 suppressors permitted a significant increase in the titer of phage VIB04 (**Table 1)**. Mutations in *pcnB* that reduced the copy number of pWR1566 displayed low phage titers that were similar to WT indicating that PcnB mutations did not interfere with the ability of VSP-2 gene products to block VIB04 replication. In contrast, strains with a single missense mutation present in *priA* permitted titers of phage as high as those seen in Δ*VSP*-2. Although PriA is also required for the replication of plasmids possessing a ColE1 origin of replication ^40^ the ability of a PriA mutation to alleviate VSP-2-dependent phage restriction suggests it plays a more critical role in CDD. The phage titers for strains carrying frameshift *ddmC Asp523Tyr* and *Phe571Cys* missense mutations are significantly more permissive for phage replication than most other *ddmABC* mutations. This indicated that many of the plasmid-selected mutations including premature stop or frameshifts were likely not completely inactivating their respective proteins. However, mutations in the DdmC C-terminus appear to be the most inhibitory regarding both plasmid and phage restriction. Thus, components of the DdmABC system together with PriA are likely critical components of the CDD system but some protein domains identified in our plasmid-based selection appeared to be more essential for phage restriction.

Orthologues of the PriA protein in other bacteria recognize stalled replication forks and DNA hairpin sequences during replication restart ^45–48^. The *priA M462R* CDD suppressor mutation we isolated is positioned within a highly-conserved domain that contains a β-hairpin/strand-separation ‘pin element’ that is located in a zinc-binding pocket that is conserved in RecQ helicases ^48^ (**Figure 2a**). Mutations within this pin element reportedly eliminate PriA-mediated DNA unwinding, function, and interactions with primosome protein PriB ^48^. We examined the function of the *priA M462R* in VSP-2 phage restriction by comparing the titers of phage plated on lawns of *V. cholerae* strain carrying the *priA M462R* mutation (**Table 1**). The higher titers of VIB04 on these mutants compared to the low titers on WT support the importance of PriA and its β-hairpin/strand-separation pin element in phage restriction by the CDD system and suggest that PriA may directly recognize critical DNA structures or protein complexes triggered by hairpin sequences. We were unable to construct complete *priA* deletions mutants to compare to the *priA M462R*. In *E*.*coli, priA* disruption mutants are constitutively SOS-induced, have poor viability, and form filamentous cells ^49–51^. Because *E*.*coli priA* null mutants are unable to maintain ColE1 replicon plasmids ^40^, and we measured a reduction in plasmid copy number for this mutant in *V. cholerae*, we hypothesize *priA M462R* is impaired in its activity rather than inactive in *V. cholerae*.

### Mutations in a hairpin encoding palindromic sequence abrogate CDD

To more precisely examine the role of palindromic sequences that might encode ssDNA hairpins, we inserted a minimal short synthetic oligonucleotide that encodes an RNAse III-like sequence derived from VIBO4 phage into a vector devoid of a T7 RNase III sites. This VIBO4 palindrome was designated Seq2 and was compared with an oligonucleotide that was mutated at several nucleotides predicted to disrupt base pair interactions that could drive hairpin formation (designated Seq2-mut3). The two cloned palindromic sequences were then used to evaluate CDD in the context of VSP-2 (**Figure 4abcd**). Plasmid constructs were transformed into cells, grown to stationary phase in broth, and then plated at high density on agar for 12 hrs; this is our standard CDD assay in all work reported here. We found that the Seq2 sequence was highly toxic in the CDD assay and comparable to the original T7 RNase-III sequence we identified in pWR1566. In contrast, the Seq2-mut3 encoding plasmid was ∼25-fold more permissive for the recovery of viable cells; this was comparable to the improved viability of Δ*VSP-2* strains on agar that carry a Seq2-encoding plasmid (**Figure 4d**). The phage-encoded Seq2 was just as toxic when cloned in forward and reverse orientations suggesting it is not mRNA hairpins that are triggering CDD. We conclude that one CDD signal must include palindromic sequences that can form hairpins in ssDNA or disruption of DNA replication.

**Figure 4.**
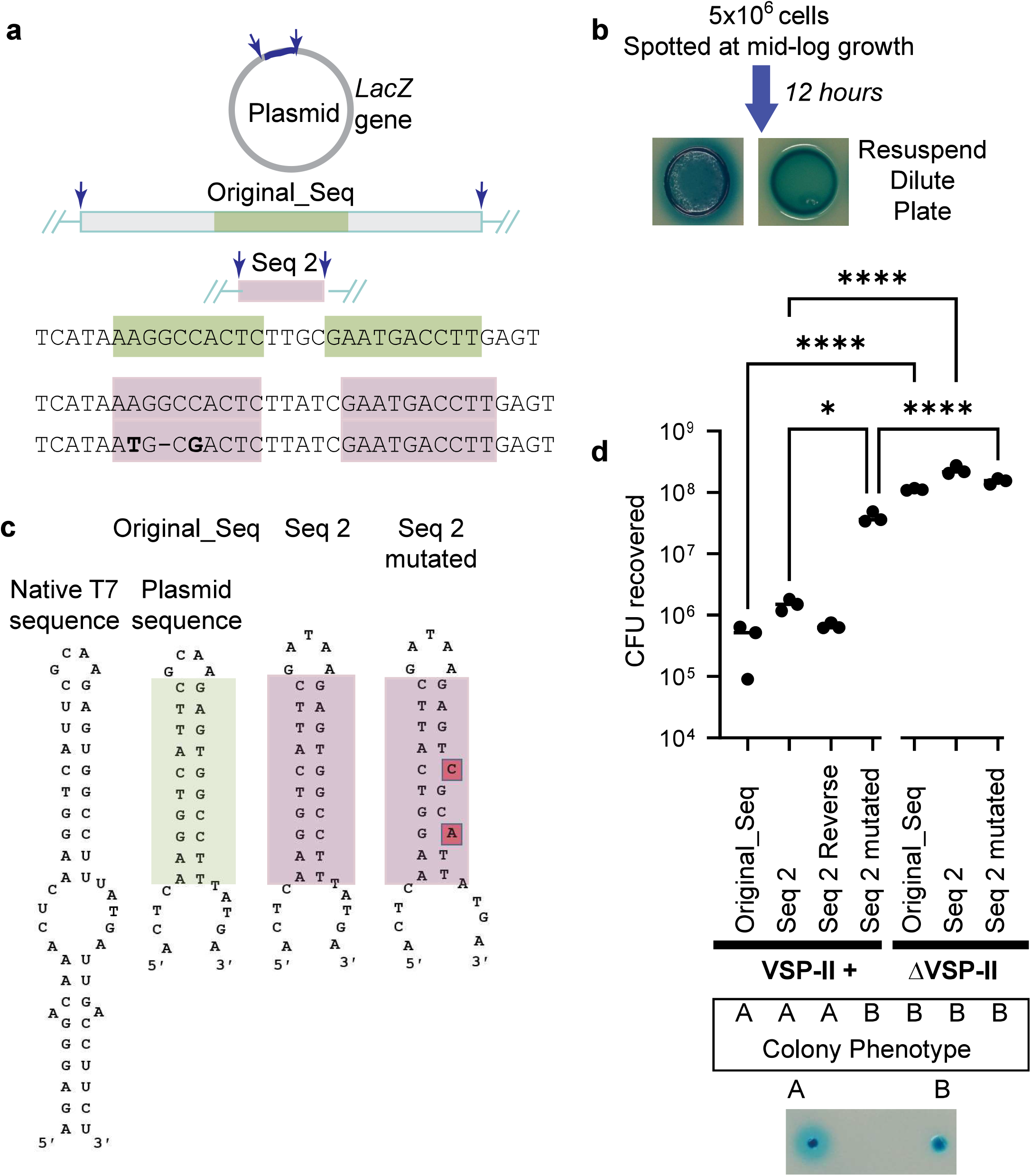
Mutations that disrupt the hairpin sequence eliminate cell-dependent death. a|A Schematic showing the position of the original sequence and its replacement by a synthetically designed sequence seq 2. The hairpin-forming nucleotides in both the original and seq2 are boxed in green and purple. b|The predicted hairpin of the original and seq2 sequences and its derivatives are shown. Mutations are boxed to indicate potential disruptions to hairpin formation. c| Screening assay showing the presence or absence of CDD after 12 hours. d| Cells recovered after 12 hours for each plasmid in both WT and Δ*VSP-2* mutants. The colony morphology of plated transformed cells is scored for each. The “blue ghost” phenotype is indicated by “A”. Normal blue colonies are indicated by “B”. Significance is indicated (* < 0.05 and **** <0.0001).

### CDD causes phage restriction and abortive infection

We characterized the role of VSP-2 in restricting the growth of vibriophage VIB04 in both high-density liquid cultures and when spotted on agar surfaces. We measured phage adsorption and growth in WT and a Δ*VSP-2* strain to establish phage infectivity growth parameters as well as the effects of VSP-2 on cell viability. When both strains are infected in liquid culture at a low multiplicity of infection but high cell density, only the Δ*VSP-2* mutant strain continues to produce infectious particles **(Figure 5a)**. In contrast, WT strains incubated with phage continue to reduce the number of infectious particles over time. After 150 minutes, Δ*VSP-2* had nearly 100,000-fold more PFUs present in supernatant fluids than WT. Cells carrying VSP-2 failed to produce an increase in infectious phage when compared to the Δ*VSP-2* strain.

**Figure 5.**
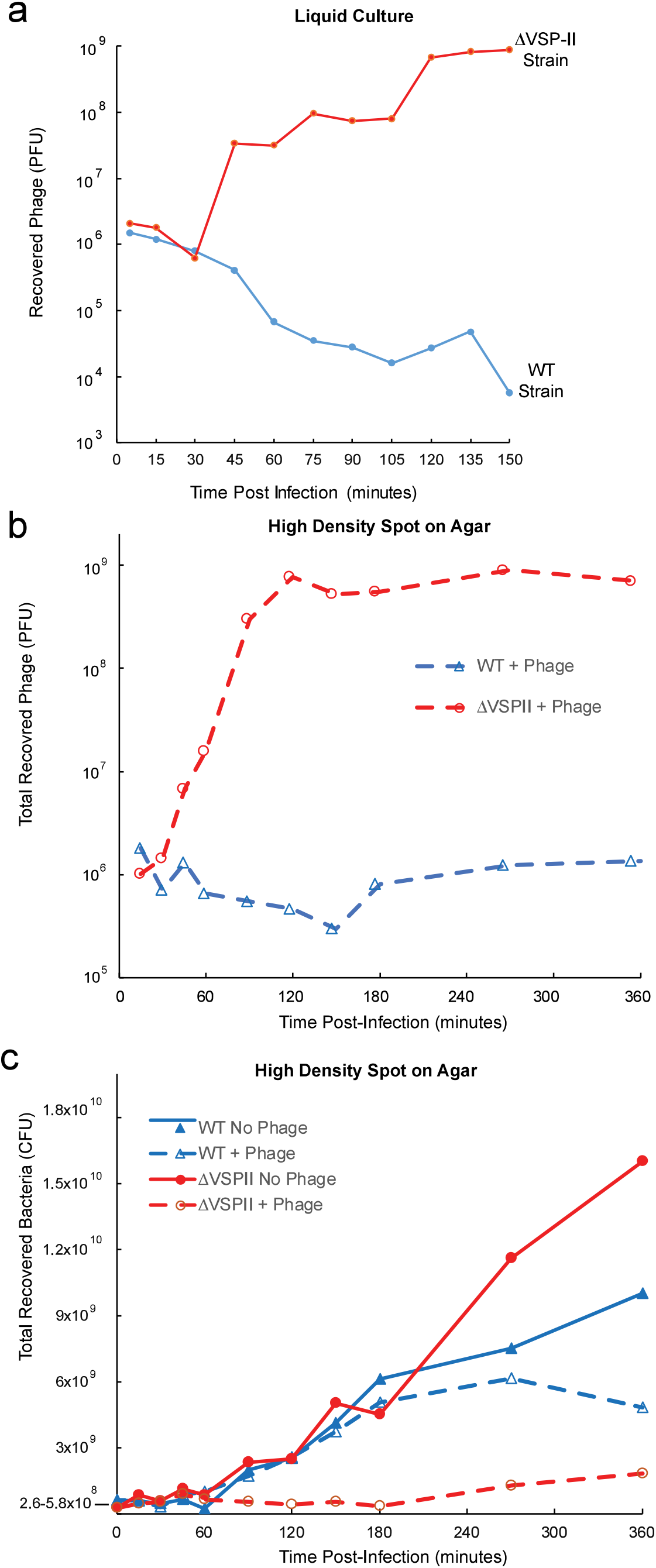
Phage production and cell viability during phage infection are impacted by VSP-II. a |Plaque forming units (PFU) representing the production of infectious phage in both WT and DVSP-II mutant strains in liquid culture for 150 minutes. b and c| Cells were infected with VIB04 phage and spotted at high density on agar to mimic CCD conditions. Both phage (PFU) and viable bacterial (CFU) were enumerated for six hours post-infection.

We surmised WT cells are also infected and fail to produce phage but did not know whether infected cells are killed by an abortive infection mechanism or instead only restrict phage growth but remain viable. To simultaneously measure the production of phage (PFUs) and viability of bacteria (CFUs), we infected cells to allow phage to adsorb and then collected the cells and plated these as a concentrated spot to mimic conditions that induced plasmid-dependent CDD. Relative to the wild type, the Δ*VSP-2* mutant grew to a higher density after 4-6 hours in the absence of phage, but CFU recovery was modestly reduced when phage-infected (**Figure 5b)**. The difference in the production of phage indicated by an increase in PFU was highly significant between WT and the Δ*VSP-2* mutant (**Figure 5c)**. Consistent with our measurements in liquid growth, only *V. cholerae* Δ*VSP-2* mutants are successfully infected and continue to produce infectious particles. At high plating density, both the number of WT and Δ*VSP-2* mutant cells were found to decrease. Thus, it is likely that phage infection, though DdmABC-restricted, can induce cell death of the WT strain through DDC abortive infection in contrast to killing the Δ*VSP-2* mutant through viral replication in these phage-permissive cells.

### Sensitivity to DNA damaging agents

Our data indicate that the *cis-*acting palindromic sequences in both phage and plasmid DNA are lethal signals to bacteria in both a DdmABC and PriA-dependent manner. How cell death is triggered likely involves activation of the CAP4 nuclease CAP4 domain in DdmA because a defective nuclease mutant was previously demonstrated to abolish plasmid elimination that depended on *ddmABC*^8^. The significant hairpin-forming structures identified as CDD-inducing signals may stall replication forks that then recruit PriA as suggested in earlier studies ^45,48,52^. We hypothesized that DNA damage may induce similar DNA repair responses that require PriA and could trigger CDD. Therefore we tested WT and CDD-defective strains for differences in their sensitivity to various mutagens and bactericidal agents that target DNA replication. We found that at a range of ultraviolet radiation doses, both single and double Δ*VSP-2* and *priA Met462Arg* mutants showed much better growth and survival than their WT parental strain (**Figure 6**). At doses above 48 joules/cm2, only strains with either or both Δ*VSP-2* and *priA* mutations were recovered. We measured the sensitivity of Δ*VSP-2* and *priA* mutants to the DNA alkylating agent methyl methane sulfonate (MMS), nalidixic acid (which reversibly binds and inhibits both the ligation activity of DNA gyrase and Topoisomerase IV resulting in dsDNA breaks^53,54^), and Mitomycin C (which introduces crosslinks between bases and dsDNA breaks that require PriA-dependent repair ^55,56^). When compared to Δ*VSP-2* and *priA* mutants, the WT strain was significantly more sensitive to the lethal actions of MMS, nalidixic acid, and Mitomycin C (**Figure 6**). Thus the same gene products that are essential for CDD also hyper-sensitize *V. cholerae* to agents that cause DNA damage.

**Figure 6.**
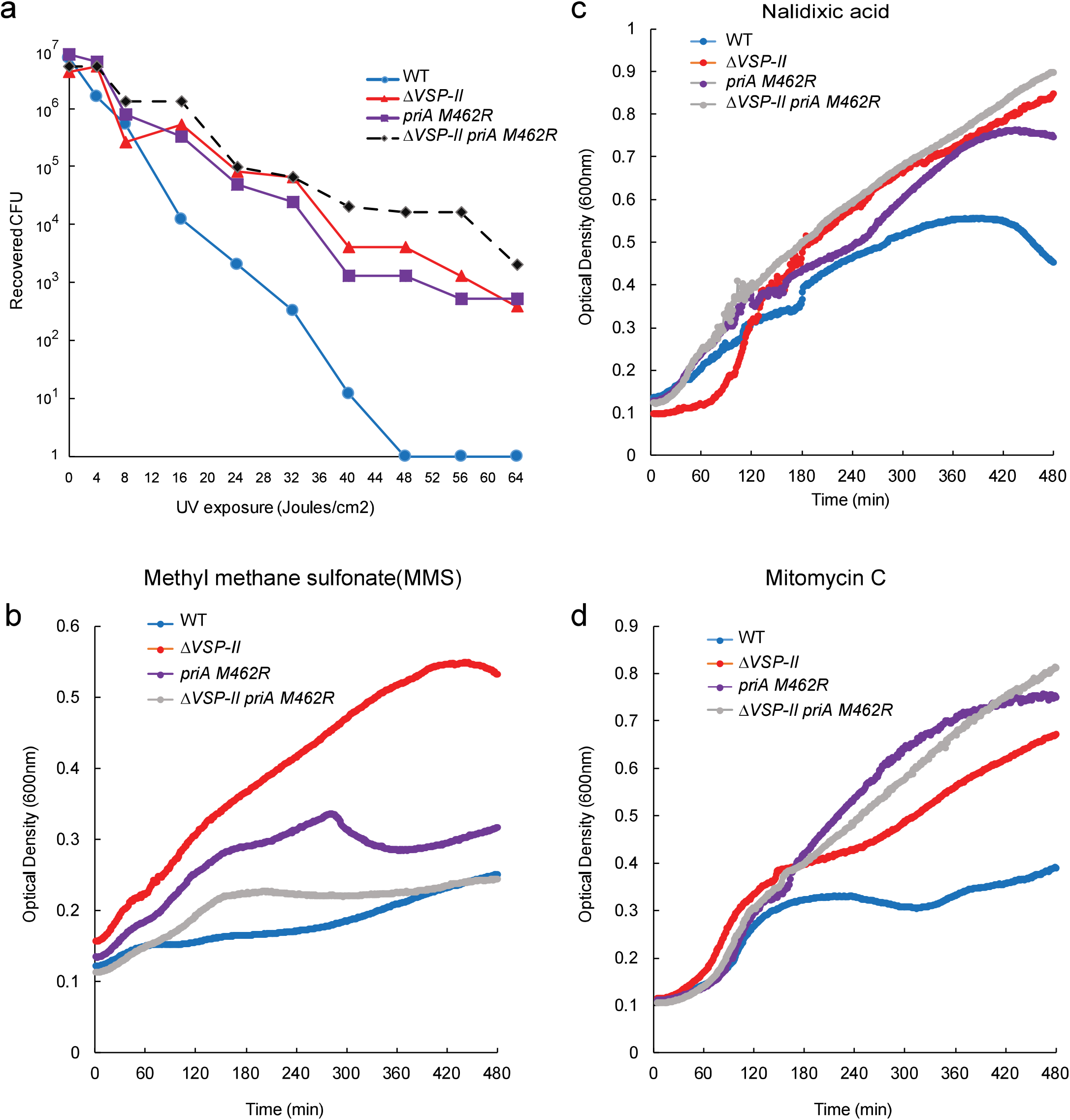
Sensitivity to UV and DNA-damaging chemicals is enhanced by VSP-II. a |Measured doses of UV exposure between 0 and 64 Joules/cm^2^ to spotted cells on agar reveal the sensitivity based on recovered CFU. b-d |Comparison of subinhibitory concentrations of Methyl methane sulfonate (MMS) 30ug/ml, Nalidixic acid (Nal) 50ng/ml, and Mitomycin C 5ng/ml (MC) to the growth of WT and both D*vsp-II* and *priA* mutants in liquid culture.

## Discussion

We show the Lamassu system of *V. cholerae* encoded by *DdmABC* of VSP-2 genomic island restricts some vibriophages by recognizing a signal associated with phage-encoded palindromic DNA sequences. This signal-induced cell density-dependent death (CDD) is dependent on the quorum-regulated *DdmABC* operon of VSP-2 ^36^. The signal was first identified in a laboratory plasmid vector as a coliphage T7-derived phage sequence. However, palindromic sequences present in two different vibriophages were also recognized by the gene products we identified as being required for VSP-2 phage restriction. CDD triggered by one of these palindromic sequences could be abolished by only three nucleotide substitutions that are predicted to disrupt the DNA hairpin it can form.

Based on these predictions, we hypothesize the DdmA nuclease effector is activated by a signal recognized in part by its DdmC and DdmB partner proteins and also possibly PriA. Critically, proteins that resemble DdmC and other structural maintenance of chromosome (SMC) proteins often play essential roles in DNA tethering, repair, replication, and recombination, usually as dimers associated with larger multimeric complexes ^57,58^. SMC-like proteins possess ATP and nucleotide-binding domains as well as an extended coil-coil domain capped by a dimerization domain ^59–61^. One of the best studied, DNA repair protein Rad50, also forms MRe11/NSB1 complexes that respond to viral invasion in eukaryotic cells and specifically recognize DNA double-stranded breaks (DSBs) and stalled replication forks ^62^. SMC bacterial repair protein SbcC forms a complex with its nuclease partner SbcD to recognize and cleave palindromic sequences that can form hairpins during DNA replication ^63,64^. Palindromic sites cause genomic instability and have been shown to induce single and double-stranded breaks via SbcCD recognition; DSB repair typically requires RecBCD recombination and the re-established replication fork protein PriA ^65^. Thus, the identified suppressors mapping to DdmABC and PriA suggest that the signal that triggers CDD may involve DNA lesions, breaks, and stalled replication forks induced by palindromic sequences. Although DdmABC proteins can resemble others that help coordinate DNA repair, recombination, replication, and maintain chromosomal stability, our results suggest that the *V. cholerae* DdmABC/PriA system instead acts to block replication of foreign DNA by recognizing DNA damage induced by these foreign genetic elements. The foreign DNA is likely not selectively destroyed but rather CDD triggers an abortive infection that eliminates both the infected cell and the viral or plasmid element that has invaded the cell before it has had a chance to amplify and infect other cells. This is most vividly demonstrated by the profound bacterial cell death that occurs when cells carrying plasmids that encode phage-derived palindromic sequences are plated at high density on agar. (**Figure 1**).

The *priA Met462Arg* allele renders cells equally permissive for phage growth as those deleted in *ddmABC* and the mutations together do not increase the plating efficiency. Curiously, PriA has been shown to recognize stalled replication complexes generated when palindromic sequences interfere with DNA replication processes ^48^. PriA is universally conserved in bacteria and likely works with DdmABC proteins to recognize PriA-DNA complexes that are uniquely formed by either palindromic sequences or certain types of DNA damage (**Figure 7**). For bacteria such as *V. cholerae* that encounter phage in many environments and the host, the DdmABC system may have a profound influence on phage infection as well as the acquisition of at least some plasmids. A recent finding reveals that some plasmids are not stable in *V. cholerae* and that more than one abortive system may be restrictive ^8,36^. Here we have shown the DdmABC system is also highly endogenously active against certain phages that are abundant in many environments and reveal both a *cis-*acting signal and a co-requirement of the host factor PriA as minimal requirements. Furthermore, because PriA is recruited in key DNA repair and replication processes, DdmABC proteins may reduce the fitness of bacteria that encounter various stresses that damage DNA and elicit such responses. We discovered that the DdmABC system increases sensitivity to both ultraviolet radiation and chemical agents that are previously demonstrated to induce damage that recruits PriA during DNA repair ^66–68^. Thus, this phage resistance mechanism in *V. cholerae* 7^th^ pandemic strains may impose a fitness cost for environmental strains of *V. cholerae* (which typically lack the DdmABC system) if these strains experience strong ambient sunlight and UV flux or exposure to mutagenic molecules in aquatic habitats or other environments.

**Figure 7.**
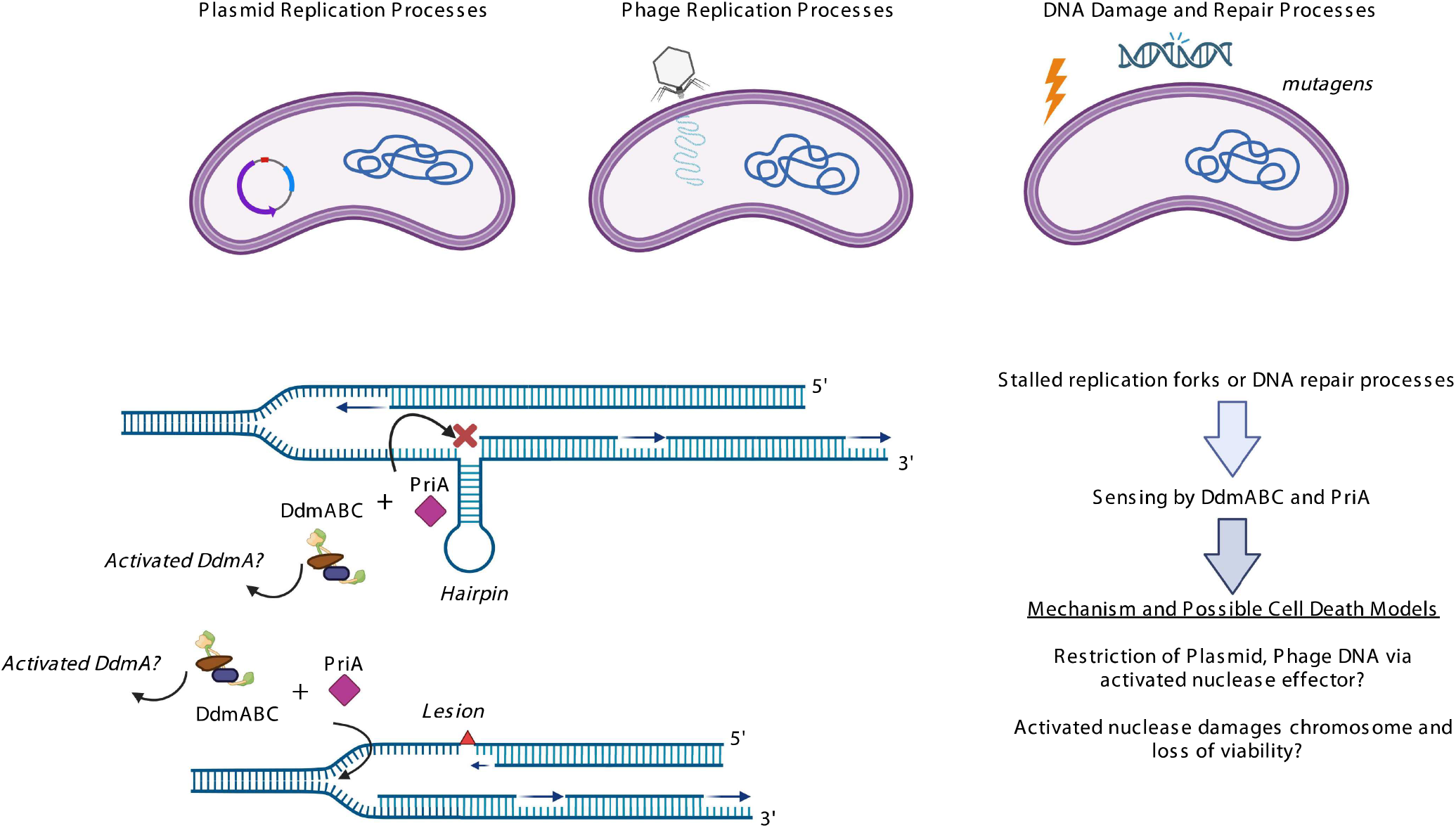
A general model of signal recognition by plasmids, phages, and DNA damage.

Recently published mechanistic work on antiphage defense systems continues to reveal striking similarities between bacteriophage abortive infection mechanisms and eukaryotic innate immunity processes ^2,3,41^. Previous work indicates RAD50-Mre11-Nbs1 DNA damage machinery componen*ts* (which are orthologous to DdmA and DdmC) are regulated and disrupted by several adenovirus-encoded proteins during infection and this is critical for maintaining viral DNA replication ^69,70^. Both Rad50 and Mre11 are implicated in the recognition of cytoplasmic dsDNA and activation of the innate immune genes including STING and IRF3 ^71^. Rad50-CARD9 signaling complexes in dendritic cells have been demonstrated to activate NF-κβ and the production of IL-1β when a dsDNA vector was transfected or when dendritic cells were infected with a DNA virus ^72^. Thus, insights gained on the mechanism of phage recognition through the DdmABC/PriA system may be highly relevant to the innate immune recognition of eukaryotic DNA viruses or the use of DNA vectors in human gene therapy.

Of note, DdmABC also possesses similarities to studied antiphage defense systems encoded on bacteriophages. The Old (*o*vercoming *l*ysogenization *d*efect) protein encoded by bacteriophage P2 kills *E*.*coli recB* and *recC* mutants and interferes with phage lambda growth by killing cells unless the lambda-encoded recombination genes are deleted ^73^. The CTD structure of Old protein also bears resemblance to Rad50 protein but possesses a fused nuclease domain similar to Mre11 at its N-terminus ^74^. Thus, even phages apparently acquired and adapted Rad50/SbcC-like proteins to sense recombination-based signals and then utilize those signals to degrade the chromosome of the host bacterium. We propose Rad50 and many other SMC**-**like proteins, that were previously classified as first responders for DNA damage recognition and repair, are also likely to be major sensors of damaged DNA replication processes that represent pathogen-associated molecular patterns (PAMPs) for the replication of foreign DNA. How this anti-phage and anti-plasmid system has shaped the evolution of *V. cholerae* is an important consideration in future studies focused on understanding the emergence of this highly successful pandemic pathogen. These insights may also be important to recognizing how mammalian cells respond to DNA viruses or vectors used in therapeutic applications.

## Methods

### Bacterial strains and plasmids

Strains and plasmids are listed in **Supplemental Table S1**. Strain TND0652 was a generous gift from Ankur Dalia and published elsewhere^75^. WPR2700 was first constructed to test the functionality of the *V*.*cholerae* O395 CRISPR-CasI-E system in a derivative of the El Tor strain E7946. To construct WPR2700, we first inserted a 1.4kb DNA fragment containing the *LacI* gene and an inducible tac promoter in front of the CRISPR-Cas complex in O395 at nt position 2827729 using suicide vector pRE107 ^76^ and 500 nts of flanking sequence. The DNA from O395 was prepared and transformed and integrated into TND0625 and we selected for uptake using ampicillin (100ug/ml). In this strain, the CRISPR-CAS-I-E merodiploid sequence was crossed out using sucrose counter-selection as previously described ^76^. Insertion of LacIq-ptac-CRISPR/Cas-IE was confirmed by PCR and sequencing. A 37,850 nucleotide region from O395 from *PurD* (Chromosome I position 311908) through *Fis* genes (Chromosome I position 349559) replaced the 20,417 nucleotide region E7946 derivative this introducing the entire CRISPR-CAS-I-E operon and several adjacent genes both up and downstream from O395. pWR1566 is derived from pUC18-miniTn7-LacZ and has a deletion of the Gentamycin resistance gene between flanking FRT sites generated by co-transformation with the pFRT2 vector expressing flip recombinase ^77^.

### CCD assay and selection and sequencing of suppressors

pWR1566 was purified from laboratory *E*.*coli* strain NEB 10-beta (New England Biolabs) and used to transform *V*.*cholerae* strain WR2700 using electroporation. Transformants were selected on LB agar supplemented with Carbenicillin (100µg/ml) and XGAL (150µg/ml). Colonies were resuspended in liquid no more than 12 hours after plating and grown in LB media supplemented with Carbenicillin (100ug/ml) at 37°C.

For screening of CCD and selection of survival mutants, transformed strains were grown to high OD600 (> 1.5) in shaking tubes or flasks at 37°C (RPM=300). 1.0ml of the culture was centrifuged at 3000 RFC for 4 minutes in a microcentrifuge. The supernatant was decanted and cells were resuspended in 100ul LB. This was serially diluted 4-fold 12 times in a 96-well plate row. 3ul of each dilution was spotted on LB agar (Carb 100ul/ml +XGal150ug/ml) and allowed to dry for 10 minutes before growth at 30°C. Growth on plates was observed 16-20 hours post-plating.

For the selection of survivors from CDD, we plugged spotted agar areas where cell growth was apparently absent. These plugs were resuspended in 1.0 ml LB, vortexed, and plated on LB agar (Carb 100ul/ml +XGal150ug/ml). Colonies that grew were scored for blue/white and purified on LB (Carb 100ul/ml +XGal150ug/ml). Plasmid and chromosomal DNA were prepared using the Zymo Genomic DNA Isolation kit (Zymo). Mutants were sequenced using Illumina Miseq and libraries were prepared using the DNA Ultra II library preparation kit (New England Biolabs). Mutations and depth were mapped using CLC-Bio Workbench v8 ^78^.

### Phage DNA shotgun cloning

Viral DNA from VIB04 was isolated from CsCl-purified particles dialyzed into 10mM Tris-Cl pH8.0 1mM EDTA and adjusted to 500ul using 10mM Tris-Cl pH8.01mM EDTA using one phenol extracted (200ul, equilibrated with 50mM Tris-Cl pH8.0). This was briefly mixed using a vortex and the top layer was carefully removed and kept after separation in a 55°C heat block for 20-30 minutes. Three successive Phenol:Chloroform: Isoamyl (20% volume, 25:24:1, phenol equilibrated with 50mM Tris-Cl pH8.0) extractions were performed as above. DNA was precipitated using three volumes of 100% ethanol and the pellet was washed twice with 80% ethanol and dried. DNA was dissolved in water, and sheared to ∼500nt using a (QSONICA Q800R3) for five minutes (30 seconds cycled ON/OFF at 20%). DNA was end-repaired using DNA Repair Mix (NEB E6050L) and shotgun cloned into pWR2700 that had been cut with SmaI (NEB R0141L) and AvrII (NEB R0174L), blunted using DNA Repair Mix, and dephosphorylated using Antarctic Phosphatase (NEB M0289L) using T4 Blunt T/A DNA ligase (NEB M0367L). Transformants were cultured and screened for CDD as described in this work.

### Cloning of artificial hairpin sequences

Complementary oligonucleotides that form seq2 and seq2_mutated were ordered. Cloned hairpin sequences are flanked by PmlI and SmaI restriction sites (**Supplemental Table S1**). Oligonucleotides were mixed in 100ul at a concentration of 5uM and heated to 95°C and slowly cooled until 50°C to anneal. DNA was digested with PmlI and SmaI (New England Biolabs), ligated into pWR2700 that had been cut with SmaI (NEB R0141L) and AvrII (NEB R0174L), blunted using DNA Repair Mix, and dephosphorylated using Antarctic Phosphatase (NEB M0289L) using T4 Blunt T/A DNA ligase (NEB M0367L). Transformants were cultured and screened for CDD as described in this work.

### Phage screening, preparation, and infection assays

A panel of *V*.*cholerae* phage lysates was prepared on *V*.*cholerae* strain MAK757 and plated on both WR2700 and WR2700 *ΔVSP-2*. Two candidate phages were identified as VSP-2 restricted. Both phages were sequenced and one was previously identified as Figure *V*.*cholerae* phage N4. To avoid confusion, we renamed the N4 phage “VIB04” in this work because another unrelated phage also called N4 has been studied previously in the field. The second phage (VIB05) was isolated from lysate samples kindly provided by Dr. Shah Farque who was previously a member of the International Centre for Diarrhoeal Disease Research, Dhaka, Bangladesh.

Phages were propagated on WR2700 *ΔVSP-2*. In brief, sufficient phages were co-plated in 8ml soft top agar (0.3%) with bacteria (9×10^7^ CFU from liquid mid-log culture) over 1.5% LB agar and incubated overnight at 30°C to produce a confluent lysed lawn of *V*.*cholerae* cells. This was done in triplicate using 150mm x15mm plates to increase yields. 30ml LB was poured onto each soft agar overlay and then both soft top agar and LB were mixed using a sterile plate spreader and the slurry was poured into a centrifuge bottle. This mixture was centrifuged at 15,000 RFC for 20 minutes to remove solids. The supernatant was adjusted to 1.0M NaCl and polyethylene glycol (PEG) (MW 8000). was added to 0.8% w/v. This solution was incubated on ice for between 4 to 8 hours and then centrifuged at 8,000RFC for 30 minutes (4°C). The supernatant was poured off and the pellet was gently resuspended in 50mM Tris-Cl pH8.0 and 10mM MgCl2 and incubated on ice for 1-2 hours to suspend phage virions. This suspension was centrifuged at 5,000 RFC for 10 minutes (4°C) to remove large insoluble material and the supernatant was collected. The supernatant was overlayed onto a CsCl step gradient (ρ =1.62, 1.53, and 1.62) and spun at 40,000 RPM for 90 minutes using an SW55.1 rotor on a Beckman Ultracentrifuge. The phage band was collected between ρ=1.53 and ρ =1.62 steps and dialyzed into T7 Buffer (100mM Tris-Cl pH7.5, 1.0M NaCl, 1mM EDTA).

Phages were titered on WPR2700 *ΔVSP-2* to measure the concentration of infectious particles. For phage lysis and infected cell survival assays, phage was infected at selected MOI in liquid culture and for agar assays, infected cells were pelleted at 4,000 RFC and spot plated on 1.5 LB agar. Bacteria were resuspended in 1.0ml at selected time points and the supernatant was both titered on WPR2700Δ*VSP-2* and plated on LB agar to determine phage and bacterial concentrations.

### Cell viability and growth after UV and mutagenic exposure

To measure survival after UV exposure, 10^8^ bacterial cells from a growing late log culture (OD 1.4) were spotted on LB agar and exposed to a range of UV doses determined by time exposure settings on a UV Stratalinker. Agar with spotted cells was removed as a plug and cells were resuspended in 1.0ml LB, serially diluted, and plated on LB agar to measure CFU.

To measure the effects of mutagenic compounds on bacterial growth, strains were grown shaking rigorously in 200ul LB in a 96-well plate using a Biotek Synergy MX at 30°C. Initial screening for inhibitory concentrations of each was first determined using a range of serial dilutions. A concentration that was minimally inhibitory for WPR2700 was chosen and used for the entire panel of strains. When growth curves are compared, bacterial cultures were grown to mid-log (OD 1.0), diluted 10X, and added to wells supplemented with various concentrations of Nalidixic acid, Methyl Methanosulfate, or Mitomycin C. Optical density at 600nm was measured during cell growth at 30°C for 6 hours.

### DNA folding prediction

Predictive folding of ssDNA was completed using ViennaRNA v2.5.18 ^79^. Fold algorithm options included minimum free energy and partition function to calculate the base pairing matrix in addition to the structure. Energy parameters were set for DNA with dangling energies on both sides of a helix in any case.

### Protein structure prediction

Protein structures for DdmABC and PriA were predicted using ColabFold run locally on the Orchestra cluster at Harvard Medical School ^43^. Structures were visualized using Pymol (v2.3.4)^80^ and colored to specific confidence. Proteins identified as similar to DdmC were identified using HHPRED in the MPI toolkit ^81^.

### Statistics

In CCD assays that measured the toxicity of cloned hairpin sequences and were performed in triplicate, we applied an ordinary one-way in PRISM (v9.4.1) for Windows.

## References

1. Labrie, S. J., Samson, J. E. & Moineau, S. Bacteriophage resistance mechanisms. Nature Reviews Microbiology 8, 317–327 (2010).

2. Duncan-Lowey, B. & Kranzusch, P. J. CBASS phage defense and evolution of antiviral nucleotide signaling. Current Opinion in Immunology 74, 156–163 (2022).

3. Wein, T. & Sorek, R. Bacterial origins of human cell-autonomous innate immune mechanisms. Nature Reviews Immunology 22, 629–638 (2022).

4. Doron, S. et al. Systematic discovery of antiphage defense systems in the microbial pangenome. Science 359, eaar4120 (2018).

5. Makarova, K. S., Wolf, Y. I. & Koonin, E. V. Comparative genomics of defense systems in archaea and bacteria. Nucleic Acids Research 41, 4360–4377 (2013).

6. Whiteley, A. T. et al. Bacterial cGAS-like enzymes synthesize diverse nucleotide signals. Nature 567, 194–199 (2019).

7. Cohen, D. et al. Cyclic GMP–AMP signalling protects bacteria against viral infection. Nature 574, 691–695 (2019).

8. Jaskólska, M., Adams, D. W. & Blokesch, M. Two defence systems eliminate plasmids from seventh pandemic Vibrio cholerae. Nature 604, 323–329 (2022).

9. Comeau, A. M. et al. Exploring the prokaryotic virosphere. Research in Microbiology 159, 306–313 (2008).

10. Clokie, M. R. J., Millard, A. D., Letarov, A. V. & Heaphy, S. Phages in nature. Bacteriophage 1, 31–45 (2011).

11. Stern, A. & Sorek, R. The phage-host arms race: Shaping the evolution of microbes. BioEssays 33, 43–51 (2011).

12. Hampton, H. G., Watson, B. N. J. & Fineran, P. C. The arms race between bacteria and their phage foes. Nature 577, 327–336 (2020).

13. Lander, E. S. The Heroes of CRISPR. Cell 164, 18–28 (2016).

14. Salmond, G. P. C. & Fineran, P. C. A century of the phage: past, present and future. Nature Reviews Microbiology 13, 777–786 (2015).

15. Kaper, J. B., Morris, J. G. & Levine, M. M. Cholera. Clin. Microbiol. Rev. 8, 48 (1995).

16. Faruque, S. M. & Mekalanos, J. J. Pathogenicity islands and phages in Vibrio cholerae evolution. Trends in Microbiology 11, 505–510 (2003).

17. Waldor, M. K. & Mekalanos, J. J. Lysogenic Conversion by a Filamentous Phage Encoding Cholera Toxin. Science 272, 1910 (1996).

18. Taylor, R. K., Miller, V. L., Furlong, D. B. & Mekalanos, J. J. Use of phoA gene fusions to identify a pilus colonization factor coordinately regulated with cholera toxin. Proc Natl Acad Sci USA 84, 2833 (1987).

19. Herrington, D. A. et al. Toxin, toxin-coregulated pili, and the toxR regulon are essential for Vibrio cholerae pathogenesis in humans. Journal of Experimental Medicine 168, 1487–1492 (1988).

20. Hassan, F., Kamruzzaman, M., Mekalanos, J. J. & Faruque, S. M. Satellite phage TLCϕ enables toxigenic conversion by CTX phage through dif site alteration. Nature 467, 982–985 (2010).

21. Faruque, S. M., Albert, M. J. & Mekalanos, J. J. Epidemiology, Genetics, and Ecology of Toxigenic Vibrio cholerae. Microbiol. Mol. Biol. Rev. 62, 1301 (1998).

22. Mutreja, A. et al. Evidence for several waves of global transmission in the seventh cholera pandemic. Nature 477, 462–465 (2011).

23. Domman, D. et al. Integrated view of Vibrio cholerae in the Americas. Science 358, 789 (2017).

24. Weill, F.-X. et al. Genomic history of the seventh pandemic of cholera in Africa. Science 358, 785 (2017).

25. Ramamurthy, T. et al. Virulence patterns of Vibrio cholerae non-01 strains isolated from hospitalised patients with acute diarrhoea in Calcutta, India. Journal of Medical Microbiology vol. 39 310–317 (1993).

26. Sharma Charu et al. Molecular Analysis of Non-O1, Non-O139 Vibrio cholerae Associated with an Unusual Upsurge in the Incidence of Cholera-Like Disease in Calcutta, India. Journal of Clinical Microbiology 36, 756–763 (1998).

27. Chakraborty Soumen et al. Virulence Genes in Environmental Strains of Vibrio cholerae. Applied and Environmental Microbiology 66, 4022–4028 (2000).

28. Dziejman, M. et al. Comparative genomic analysis of Vibrio cholerae: Genes that correlate with cholera endemic and pandemic disease. Proc Natl Acad Sci USA 99, 1556 (2002).

29. Faruque, S. M. et al. Seasonal epidemics of cholera inversely correlate with the prevalence of environmental cholera phages. Proceedings of the National Academy of Sciences 102, 1702–1707 (2005).

30. Faruque, S. M. et al. Self-limiting nature of seasonal cholera epidemics: Role of host-mediated amplification of phage. Proceedings of the National Academy of Sciences 102, 6119–6124 (2005).

31. Nelson, E. J., Harris, J. B., Glenn Morris, J., Calderwood, S. B. & Camilli, A. Cholera transmission: the host, pathogen, and bacteriophage dynamic. Nature Reviews Microbiology 7, 693–702 (2009).

32. Seed, K. D. et al. Evidence of a Dominant Lineage Vibrio cholerae-Specific Lytic Bacteriophages Shed by Cholera Patients over a 10-Year Period in Dhaka, Bangladesh. mBio 2, e00334–10 (2011).

33. Faruque, S. M. & Mekalanos, J. J. Phage-bacterial interactions in the evolution of toxigenic Vibrio cholerae. Virulence 3, 556–565 (2012).

34. Yen, M. & Camilli, A. Mechanisms of the evolutionary arms race between Vibrio cholerae and Vibriophage clinical isolates. Int Microbiol 20, 116–120 (2017).

35. Hoque M.M., Naser I.B., Bari S.M., Zhu J., Mekalanos J.J. & Faruque S.M. Quorum Regulated Resistance of Vibrio cholerae against Environmental Bacteriophages. Sci Rep 6, 37956–37956 (2016).

36. O’Hara, B. J., Alam, M. & Ng, W.-L. The Vibrio cholerae Seventh Pandemic Islands act in tandem to defend against a circulating phage. PLOS Genetics 18, e1010250 (2022).

37. Brown, S., Brickman, E. R. & Beckwith, J. Blue ghosts: a new method for isolating amber mutants defective in essential genes of Escherichia coli. J. Bacteriol. 146, 422 (1981).

38. Albertini, A. M., Hofer, M., Calos, M. P. & Miller, J. H. On the formation of spontaneous deletions: The importance of short sequence homologies in the generation of large deletions. Cell 29, 319–328 (1982).

39. Court D.L., Gan J., Liang Y.H., Shaw G.X., Tropea J.E., Costantino N., Waugh D.S. & Ji X. RNase III: Genetics and function; structure and mechanism. Annu Rev Genet 47, 405–431 (2013).

40. Lee, E. H. & Kornberg, A. Replication deficiencies in priA mutants of Escherichia coli lacking the primosomal replication n’ protein. Proceedings of the National Academy of Sciences 88, 3029–3032 (1991).

41. Lowey, B. et al. CBASS Immunity Uses CARF-Related Effectors to Sense 3′–5′- and 2′–5′-Linked Cyclic Oligonucleotide Signals and Protect Bacteria from Phage Infection. Cell 182, 38–49.e17 (2020).

42. Krishnan, A., Burroughs, A. M., Iyer, L. M. & Aravind, L. Comprehensive classification of ABC ATPases and their functional radiation in nucleoprotein dynamics and biological conflict systems. Nucleic Acids Research 48, 10045–10075 (2020).

43. Mirdita M., Schütze K., Moriwaki Y., Heo L., Ovchinnikov S. & Steinegger M. ColabFold: making protein folding accessible to all. Nature Methods 19, 679–682 (2022).

44. Walker, P. J. et al. Changes to virus taxonomy and to the International Code of Virus Classification and Nomenclature ratified by the International Committee on Taxonomy of Viruses (2021). Archives of Virology 166, 2633–2648 (2021).

45. Sandler Steven J. & Marians Kenneth J. Role of PriA in Replication Fork Reactivation inEscherichia coli. Journal of Bacteriology 182, 9–13 (2000).

46. Tanaka, T., Taniyama, C., Arai, K.-I. & Masai, H. ATPase/helicase motif mutants of Escherichia coli PriA protein essential for recombination-dependent DNA replication. Genes to Cells 8, 251–261 (2003).

47. Tanaka, T., Mizukoshi, T., Sasaki, K., Kohda, D. & Masai, H. Escherichia coli PriA Protein, Two Modes of DNA Binding and Activation of ATP Hydrolysis. Journal of Biological Chemistry 282, 19917–19927 (2007).

48. Windgassen, T. A., Leroux, M., Sandler, S. J. & Keck, J. L. Function of a strand-separation pin element in the PriA DNA replication restart helicase. Journal of Biological Chemistry 294, 2801–5614 (2019).

49. Nurse, P., Zavitz, K. & Marians, K. Inactivation of the Escherichia coli priA DNA replication protein induces the SOS response. J Bacteriol 173, 6686–6693 (1991).

50. Sandler, S. J., Samra, H. S. & Clark, A. J. Differential Suppression of priA2::kan Phenotypes in Escherichia coli K-12 by Mutations in priA, lexA, and dnaC. Genetics 143, 5–13 (1996).

51. McCool, J. D. & Sandler, S. J. Effects of mutations involving cell division, recombination, and chromosome dimer resolution on a priA2∷kan mutant. Proceedings of the National Academy of Sciences 98, 8203–8210 (2001).

52. Voineagu, I., Narayanan, V., Lobachev, K. S. & Mirkin, S. M. Replication stalling at unstable inverted repeats: Interplay between DNA hairpins and fork stabilizing proteins. Proceedings of the National Academy of Sciences 105, 9936–9941 (2008).

53. Drlica, K. et al. Quinolones: Action and Resistance Updated. Current Topics in Medicinal Chemistry 9, 981–998 (2009).

54. Lundin, C. et al. Methyl methanesulfonate (MMS) produces heat-labile DNA damage but no detectable in vivo DNA double-strand breaks. Nucleic Acids Research 40, 5794–5794 (2012).

55. Tomasz, M. Mitomycin C: small, fast and deadly (but very selective). Chemistry & Biology 2, 575–579 (1995).

56. Kogoma T., Cadwell G.W., Barnard K.G., & Asai T. The DNA replication priming protein, PriA, is required for homologous recombination and double-strand break repair. Journal of Bacteriology 178, 1258–1264 (1996).

57. Graumann, P. L. & Knust, T. Dynamics of the bacterial SMC complex and SMC-like proteins involved in DNA repair. Chromosome Research 17, 265–275 (2009).

58. Uhlmann, F. SMC complexes: from DNA to chromosomes. Nature Reviews Molecular Cell Biology 17, 399–412 (2016).

59. Hirano, T. The ABCs of SMC proteins: two-armed ATPases for chromosome condensation, cohesion, and repair. Genes & Development 16, 399–414 (2002).

60. Pellegrino S., Radzimanowski J., de Sanctis D., Boeri Erba E., McSweeney S. & Timmins J. Structural and Functional Characterization of an SMC-like Protein RecN: New Insights into Double-Strand Break Repair. Structure 20, 2076–2089 (2012).

61. Schiller, C. B., Seifert, F. U., Linke-Winnebeck, C. & Hopfner, K.-P. Cold Spring Harbor Perspectives in Biology 6, (2014).

62. Syed, A. & Tainer, J. A. The MRE11–RAD50–NBS1 Complex Conducts the Orchestration of Damage Signaling and Outcomes to Stress in DNA Replication and Repair. Annu. Rev. Biochem. 87, 263–294 (2018).

63. Connelly, J. C., de Leau, E. S. & Leach, D. R. DNA cleavage and degradation by the SbcCD protein complex from Escherichia coli. Nucleic Acids Res 27, 1039–1046 (1999).

64. Connelly, J. C., de Leau, E. S. & Leach, D. R. F. Nucleolytic processing of a protein-bound DNA end by the E. coli SbcCD (MR) complex. DNA Repair 2, 795–807 (2003).

65. Eykelenboom, J. K., Blackwood, J. K., Okely, E. & Leach, D. R. F. SbcCD Causes a Double-Strand Break at a DNA Palindrome in the Escherichia coli Chromosome. Molecular Cell 29, 644–651 (2008).

66. Ivancić-Bacće, I., Vlasić, I., Cogelja-Cajo, G., Brcić-Kostić, K. & Salaj-Smic, E. Roles of PriA protein and double-strand DNA break repair functions in UV-induced restriction alleviation in Escherichia coli. Genetics 174, 2137–2149 (2006).

67. Kline, K. A. & Seifert, H. S. Mutation of the priA gene of Neisseria gonorrhoeae affects DNA transformation and DNA repair. J Bacteriol 187, 5347–5355 (2005).

68. Madison, K. E., Abdelmeguid, M. R., Jones-Foster, E. N. & Nakai, H. A New Role for Translation Initiation Factor 2 in Maintaining Genome Integrity. PLOS Genetics 8, e1002648 (2012).

69. Evans Jared D. & Hearing Patrick. Relocalization of the Mre11-Rad50-Nbs1 Complex by the Adenovirus E4 ORF3 Protein Is Required for Viral Replication. Journal of Virology 79, 6207–6215 (2005).

70. Karen Kasey A., Hoey Peter J., Young C. S. H., & Hearing Patrick. Temporal Regulation of the Mre11-Rad50-Nbs1 Complex during Adenovirus Infection. Journal of Virology 83, 4565–4573 (2009).

71. Kondo T., Kobayashi J., Saitoh T., Maruyama K., Ishii K.J., Barber G.N., Komatsu K., Akira S. & Kawai T. DNA damage sensor MRE11 recognizes cytosolic double-stranded DNA and induces type I interferon by regulating STING trafficking. Proceedings of the National Academy of Sciences 110, 2969–2974 (2013).

72. Roth S., Rottach A., Lotz-Havla A.S., Laux V., Muschaweckh A., Gersting S.W., Muntau A.C., Hopfner K.P., Jin L., Vanness K., Petrini J.H., Drexler I., Leonhardt H. & Ruland J. Rad50-CARD9 interactions link cytosolic DNA sensing to IL-1β production. Nature Immunology 15, 538–545 (2014).

73. Finkel, S., Hailing, C. & Calendar, R. Selection of lambda Spi− transducing phages using the P2 old gene cloned onto a plasmid. Gene 46, 65–69 (1986).

74. Schiltz, C. J., Lee, A., Partlow, E. A., Hosford, C. J. & Chappie, J. S. Structural characterization of Class 2 OLD family nucleases supports a two-metal catalysis mechanism for cleavage. Nucleic Acids Research 47, 9448–9463 (2019).

75. Dalia, T. N., Chlebek, J. L. & Dalia, A. B. A modular chromosomally integrated toolkit for ectopic gene expression in Vibrio cholerae. Scientific Reports 10, 15398 (2020).

76. Edwards, R. A., Keller, L. H. & Schifferli, D. M. Improved allelic exchange vectors and their use to analyze 987P fimbria gene expression. Gene 207, 149–157 (1998).

77. Choi K.H., Gaynor J.B., White K.G., Lopez C., Bosio C.M., Karkhoff-Schweizer R.R. & Schweizer H.P. A Tn7-based broad-range bacterial cloning and expression system. Nature Methods 2, 443–448 (2005).

78. CLC Genomics Workbench.

79. Lorenz R., Bernhart S.H., Höner Zu Siederdissen C., Tafer H., Flamm C., Stadler P.F. & Hofacker I.L. ViennaRNA Package 2.0. Algorithms for Molecular Biology 6, 26 (2011).

80. The PyMOL Molecular Graphics System.

81. Gabler F., Nam S.Z., Till S., Mirdita M., Steinegger M., Söding J., Lupas A.N. & Alva V. Protein Sequence Analysis Using the MPI Bioinformatics Toolkit. Current Protocols in Bioinformatics 72, e108 (2020).

