## Supplemental Files for "Cell density-dependent death triggered by viral palindromic DNA sequences"

### Supplemental Figure S1

The sequences of three CCD-suppressing deletions identified in pWR1566 isolated in WR2700.

### Supplemental Figure S2

Conceptual Model of DdmABC compared to RAD50/Mre11. Comparison of ColabFold-predicted DdmC to various SMC family proteins identified by HHPRED

### Supplemental Figure S3

The sequences of phage VIB04 sequences found to be toxic when shotgun cloned into the pWR1566 between SmaI and AvrII restriction sites.

### Supplemental Table S1

Plasmids, strains, and oligonucleotides used in this work

**Supplemental Figure 1. Sequences adjacent to deletions identified in pWR1566**

Deletion 1: 2132-2306

AGGTCATTC|TTACCCAAC

AGGTCATTC|GCAAGAGTGGCCT..AAACCCTGGCG|TTACCCAAC

Deletion 2: 1895-3759

TTCGATCCA|CGTGGATGA

TTCGATCCA|TTGCTGTTGACAA..ATGTACGCGCG|CGTGGATGA

Deletion 3: 1941-4622

CCTATTCCG|TTGGCGTAA

CCTATTCCG|AAGTTCCTATTCT..GGATAACGACA|TTGGCGTAA

**Supplemental Figure S2**

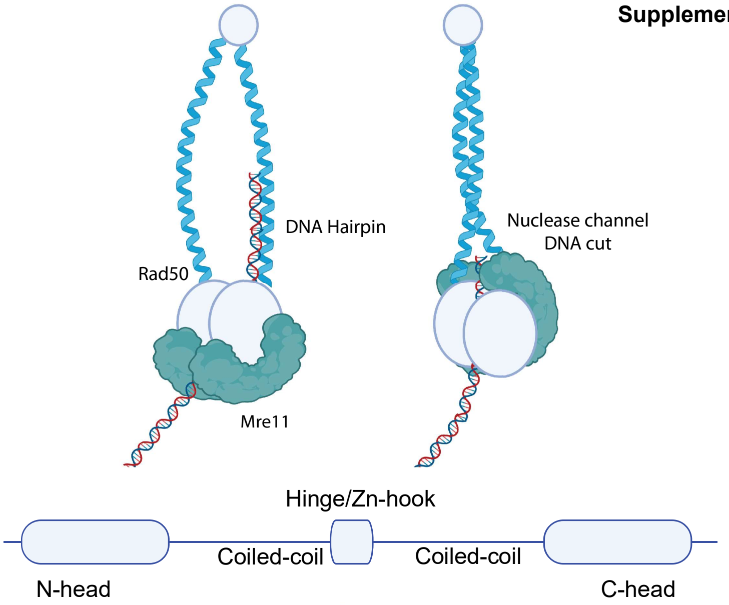


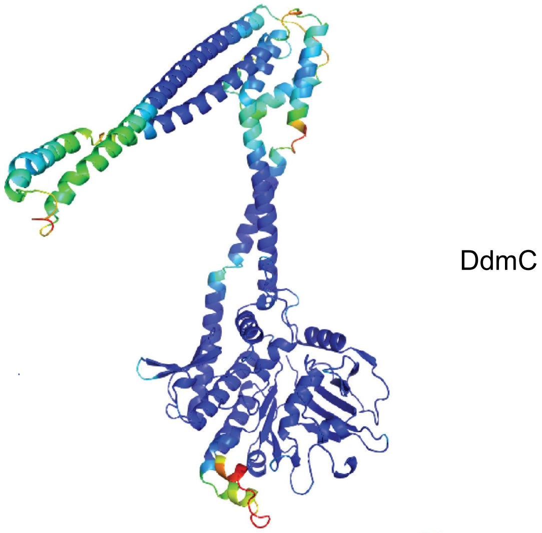


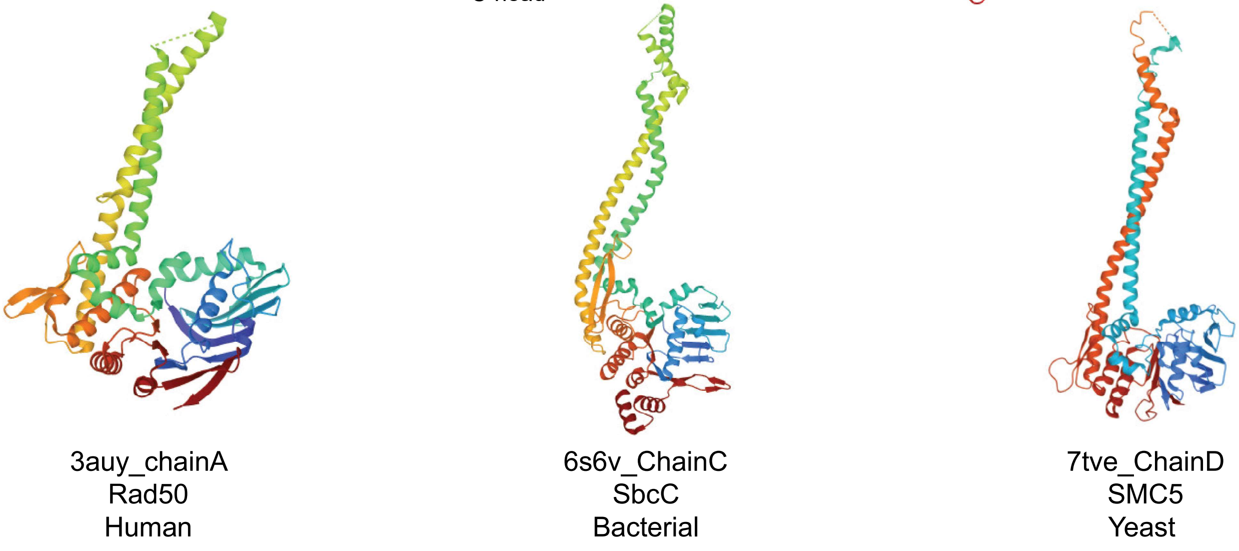

### Supplemental Figure S3

#### 118_sequence 1

```
Query  1      TGAACGTATGTTGAAATCTTCAAGTGTTACTGCTGAACAATTCGAGAACGTTAAGCAACT  60
              ||||||||||||||||||||||||||||||||||||||||||||||||||||||||||||
Sbjct  31934  TGAACGTATGTTGAAATCTTCAAGTGTTACTGCTGAACAATTCGAGAACGTTAAGCAACT  31993

Query  61     CATTCGGGACTATGCGACAGTTGATGCTAACGGATACATCAAGATTAACAACAAGTCTGC  120
              ||||||||||||||||||||||||||||||||||||||||||||||||||||||||||||
Sbjct  31994  CATTCGGGACTATGCGACAGTTGATGCTAACGGATACATCAAGATTAACAACAAGTCTGC  32053

Query  121    CTTTGCGAGTGACCCTCGTGCTATGGACTTGTGGCGTCTTGGTGATAAAATGGCTGATGA  180
              ||||||||||||||||||||||||||||||||||||||||||||||||||||||||||||
Sbjct  32054  CTTTGCGAGTGACCCTCGTGCTATGGACTTGTGGCGTCTTGGTGATAAAATGGCTGATGA  32113

Query  181    AGCTATCCTTCGTCCTACCAAAGTTTCCATGCAGAATACCAAAGCCTATGGCTCACTGGT  240
              ||||||||||||||||||||||||||||||||||||||||||||||||||||||||||||
Sbjct  32114  AGCTATCCTTCGTCCTACCAAAGTTTCCATGCAGAATACCAAAGCCTATGGCTCACTGGT  32173

Query  241    TCAACTTGGTATGCAGTTCAAATCGTTCACTCTAAAGTCTCTGAATGGTCGTACTATCCG  300
              ||||||||||||||||||||||||||||||||||||||||||||||||||||||||||||
Sbjct  32174  TCAACTTGGTATGCAGTTCAAATCGTTCACTCTAAAGTCTCTGAATGGTCGTACTATCCG  32233

Query  301    TGCAATCTATGAGGGAACTAAGAATGGACGAGCCATTGACCAGACCATTGCTGCTACATT  360
              ||||||||||||||||||||||||||||||||||||||||||||||||||||||||||||
Sbjct  32234  TGCAATCTATGAGGGAACTAAGAATGGACGAGCCATTGACCAGACCATTGCTGCTACATT  32293

Query  361    GTCTATGGGATTAGCTGCTGGTTTCTATGCGATTCGTGCTCAAGTTGCTGCTCAAGGTAT  420
              ||||||||||||||||||||||||||||||||||||||||||||||||||||||||||||
Sbjct  32294  GTCTATGGGATTAGCTGCTGGTTTCTATGCGATTCGTGCTCAAGTTGCTGCTCAAGGTAT  32353

Query  421    TCCAG  425
              |||||
Sbjct  32354  TCCAG  3235
```

#### 118_sequence 2

```
Query  78     TCATTAACCGTGATGAAGTAGAGCGTTATGCAGTGTTCTTTACAGGTTCAAACATTCGGG  137
              ||||||||||||||||||||||||||||||||||||||||||||||||||||||||||||
Sbjct  23179  TCATTAACCGTGATGAAGTAGAGCGTTATGCAGTGTTCTTTACAGGTTCAAACATTCGGG  23238

Query  138    TATTCGATTTGTTTACTGGTGATGAGAAGACTGTGAATGCTCCTAACGGTTTATCTTACG  197
              ||||||||||||||||||||||||||||||||||||||||||||||||||||||||||||
Sbjct  23239  TATTCGATTTGTTTACTGGTGATGAGAAGACTGTGAATGCTCCTAACGGTTTATCTTACG  23298

Query  198    TGACGTCTTCTAACCCTCGTAAAGACCTTCGTATGGTGACTGTTGCTGATTATACGTTCA  257
              ||||||||||||||||||||||||||||||||||||||||||||||||||||||||||||
Sbjct  23299  TGACGTCTTCTAACCCTCGTAAAGACCTTCGTATGGTGACTGTTGCTGATTATACGTTCA  23358

Query  258    TTCTTAATCGTAACGTATCGACTGCTCAAGGGACAACTAACACTCCAAGGGGACTGGCTC  317
              ||||||||||||||||||||||||||||||||||||||||||||||||||||||||||||
Sbjct  23359  TTCTTAATCGTAACGTATCGACTGCTCAAGGGACAACTAACACTCCAAGGGGACTGGCTC  23418

Query  318    CTTTTGGTCACTTTGGGTTGGTGGTTATCCGTGGTGGTCAATATGGTCGTACCTATCGAG  377
              ||||||||||||||||||||||||||||||||||||||||||||||||||||||||||||
Sbjct  23419  CTTTTGGTCACTTTGGGTTGGTGGTTATCCGTGGTGGTCAATATGGTCGTACCTATCGAG  23478

Query  378    TCAAAGTTAATGGAAGCGTAGAGGCATCCTTTGAGACTCCTTTAGGCGACCAAGTGGAAC  437
              ||||||||||||||||||||||||||||||||||||||||||||||||||||||||||||
Sbjct  23479  TCAAAGTTAATGGAAGCGTAGAGGCATCCTTTGAGACTCCTTTAGGCGACCAAGTGGAAC  23538

Query  438    ATGCAAAACAAATAGACATTGCGTATATCATTGACCAGTTAGCGGCACGTTTAATCAACA  497
              ||||||||||||||||||||||||||||||||||||||||||||||||||||||||||||
Sbjct  23539  ATGCAAAACAAATAGACATTGCGTATATCATTGACCAGTTAGCGGCACGTTTAATCAACA  23598

Query  498    GGGGATGGGCTGTAACTAAAGGTTCCGGTTATTTCTATTTCTCAAAGAGTGGTTCCGTTA  557
```

```
                ||||||||||||||||||||||||||||||||||||||||||||||||||||||||||||
Sbjct  23599  GGGGATGGGCTGTAACTAAAGGTTCCGGTTATTTCTATTTCTCAAAGAGTGGTTCCGTTA  23658

Query  558    TTATCAAATCTCTGGAAGTAGAGGATGGCTACAACGGGCAGTTGGCTTGGGGTATCATTA  617
              ||||||||||||||||||||||||||||||||||||||||||||||||||||||||||||
Sbjct  23659  TTATCAAATCTCTGGAAGTAGAGGATGGCTACAACGGGCAGTTGGCTTGGGGTATCATTA  23718

Query  618    ACGATGTTCAGAAGACAACTCAGTTACCTGTCTATGCGCCTAACAACTACATTATCCGAG  677
              ||||||||||||||||||||||||||||||||||||||||||||||||||||||||||||
Sbjct  23719  ACGATGTTCAGAAGACAACTCAGTTACCTGTCTATGCGCCTAACAACTACATTATCCGAG  23778

Query  678    TGTCTGGAGACCCTACGTTGAATCAGG  704
              |||||||||||||||||||||||||||
Sbjct  23779  TGTCTGGAGACCCTACGTTGAATCAGG  23805
```

118_16

```
Query  252    TTCGTATCGCTCAAGAAGCTATGCGTCGAGTTGGTGAACATTGGAATTTCCGGTGTCCAT  311
              ||||||||||||||||||||||||||||||||||||||||||||||||||||||||||||
Sbjct  13004  TTCGTATCGCTCAAGAAGCTATGCGTCGAGTTGGTGAACATTGGAATTTCCGGTGTCCAT  13063

Query  312    TAGACACTGAAGGGAAGATTGGAGCTAATTGGGCAATCTGTCACTAATGGTGTCTTTATG  371
              ||||||||||||||||||||||||||||||||||||||||||||||||||||||||||||
Sbjct  13064  TAGACACTGAAGGGAAGATTGGAGCTAATTGGGCAATCTGTCACTAATGGTGTCTTTATG  13123

Query  372    CAGGATATGAAGGTTGTTCGTTTCTATTTCTATCAGCGTAAACACGTAGAGTATCTTCTA  431
              ||||||||||||||||||||||||||||||||||||||||||||||||||||||||||||
Sbjct  13124  CAGGATATGAAGGTTGTTCGTTTCTATTTCTATCAGCGTAAACACGTAGAGTATCTTCTA  13183

Query  432    AAGTGGAGACACGAGATTCGTTATGGCTCTGCTAAGAAGGCTCTACAATACCGGATAGAA  491
              ||||||||||||||||||||||||||||||||||||||||||||||||||||||||||||
Sbjct  13184  AAGTGGAGACACGAGATTCGTTATGGCTCTGCTAAGAAGGCTCTACAATACCGGATAGAA  13243

Query  492    TCTCAAACTTATCTTAAAGCTATGCATGACTTAGCTAAATGATATACTCAAGGTCATTCT  551
              ||||||||||||||||||||||||||||||||||||||||||||||||||||||||||||
Sbjct  13244  TCTCAAACTTATCTTAAAGCTATGCATGACTTAGCTAAATGATATACTCAAGGTCATTCT  13303

Query  552    ATGAGAGTGGCCTTTATGAATATCGTTTATGATATCTAATTAACCCACACTATAGGGATA  611
              ||||||||||||||||||||||||||||||||||||||||||||||||||||||||||||
Sbjct  13304  ATGAGAGTGGCCTTTATGAATATCGTTTATGATATCTAATTAACCCACACTATAGGGATA  13363

Query  612    AGGGACGTAAGGTTTCTTATCTTAAAGATTAACTTAAAGAAGGAGGACAATATGTCTAAC  671
              ||||||||||||||||||||||||||||||||||||||||||||||||||||||||||||
Sbjct  13364  AGGGACGTAAGGTTTCTTATCTTAAAGATTAACTTAAAGAAGGAGGACAATATGTCTAAC  13423

Query  672    ACTCAGAAAACCACTATGACTTTCACTGGCAATCTATCAGTTGATTTCACGTTCATT  728
              |||||||||||||||||||||||||||||||||||||||||||||||||||||||||
Sbjct  13424  ACTCAGAAAACCACTATGACTTTCACTGGCAATCTATCAGTTGATTTCACGTTCATT  13480
```

### Supplemental Table S1. Bacterial strains, plasmids, and oligonucleotides

#### Strains

| Strain | Source |
|---|---|
| **TND0652** = E7946 SmR, Ptac-tfoX, pilA S67C, lacZ::lacIq, ΔVC1807::CmR | [75] |
| **O395 =** Virulent Classical Ogawa *V.cholerae* strain | Our strain collection |
| **WR2699 =** O395 [*VC395_8320* -> LacI -> ptac -> *cas*] | This work |
| **WR2700 =** TND0652 *purD-fis*::*WR2699 purD-fis* | This work |
| **WR2700 ΔVSP-2 =** WR2700 VC0490-VC0516 | This work |

| Vectors | Source |
|---|---|
| pWR1566 (pUC18mini-Tn7-LacZ ΔGmR) | This work |
| pRE107 | [77] |
| pWR1567 (phage DNA insert from VIB04) | This work |
| pWR1568 (phage DNA insert from VIB04) | This work |
| pWR1589 (phage DNA insert from VIB04) | This work |
| pWR1571 (pWR1566 Seq2) | This work |
| PWR1572 (pWR1566 Seq2 rev) | This work |
| pWR1573 (pWR1566 Seq2 mutated) | This work |

#### Oligonucleotides

Seq2 complementary oligonucleotides

AAGATGA**CACGTG**ACTCAAGGTCATTCGATAAGAGTGGCCTTTATGA**CCCGGG**ACTAATA

TATTAGT**CCCGGG**TCATAAAGGCCACTCTTATCGAATGACCTTGAGT**CACGTG**TCATCTT

Seq2_mutated complementary oligonucleotides

AAGATGA**CACGTG**ACTCAAGGTCATTCGATAAGAGTCGCATTATGA**CCCGGG**ACTAATA

TATTAGT**CCCGGG**TCATAATGCGACTCTTATCGAATGACCTTGAGT**CACGTG**TCATCTT
